# Dopaminergic neurons establish a distinctive axonal arbor with a majority of non-synaptic terminals

**DOI:** 10.1101/2020.05.11.088351

**Authors:** Charles Ducrot, Marie-Josée Bourque, Constantin V. L. Delmas, Anne-Sophie Racine, Dainelys Guadarrama Bello, Benoît Delignat-Lavaud, Matthew Domenic Lycas, Aurélie Fallon, Charlotte Michaud-Tardif, Samuel Burke Nanni, Freja Herborg, Ulrik Gether, Antonio Nanci, Hideto Takahashi, Martin Parent, Louis-Eric Trudeau

**Author notes:** **Corresponding author**: Louis-Éric Trudeau.

## Abstract

Chemical neurotransmission in the brain typically occurs through synapses, which are structurally and functionally defined as sites of close apposition between an axon terminal and a postsynaptic domain. Ultrastructural examinations of axon terminals established by monoamine neurons in the brain often failed to identify a similar tight pre- and postsynaptic coupling, giving rise to the concept of “diffuse” or “volume” transmission. Whether this results from intrinsic properties of such modulatory neurons remains undefined. Using an efficient co-culture model, we find that dopaminergic neurons establish an axonal arbor that is distinctive compared to glutamatergic or GABAergic neurons in both size and propensity of terminals to avoid direct contact with target neurons. Furthermore, while most dopaminergic varicosities express key proteins involved in exocytosis such as synaptotagmin 1, only ~20% of these are synaptic. The active zone protein bassoon was found to be enriched in a subset of dopaminergic terminals that are in proximity to a target cell. Irrespective of their structure, a majority of dopaminergic terminals were found to be active. Finally, we found that the presynaptic protein Nrxn-1α^SS4-^ and the postsynaptic protein NL-1^AB^, two major components involved in excitatory synapse formation, play a critical role in the formation of synapses by dopamine neurons. Taken together, our findings support the idea that dopamine neurons in the brain are endowed with a distinctive developmental program that leads them to adopt a fundamentally different mode of connectivity, compared to glutamatergic and GABAergic neurons involved in fast point-to-point signaling.

**SIGNIFICANCE STATEMENT:** Midbrain dopamine (DA) neurons regulate circuits controlling movement, motivation, and learning. The axonal connectivity of DA neurons is intriguing due to its hyperdense nature, with a particularly large number of release sites, most of which not adopting a classical synaptic structure. In this study, we provide new evidence highlighting the unique ability of DA neurons to establish a large and heterogeneous axonal arbor with terminals that, in striking contrast with glutamate and GABA neurons, actively avoid contact with the target cells. The majority of synaptic and non-synaptic terminals express proteins for exocytosis and are active. Finally, our finding suggests that, NL-1^A+B^ and Nrxn-1α^SS4-^, play a critical role in the formation of synapses by DA neurons.

## INTRODUCTION

Chemical synaptic neurotransmission in the central nervous system, such as found at glutamate and GABA synapses, is typically a fast and efficient process, with a delay between presynaptic action potential and postsynaptic activation of ionotropic receptors below 1-ms. In keeping with this fast kinetic, classical ultrastructural examination of synapses with electron microscopes has revealed that pre- and postsynaptic elements are typically in close juxtaposition, only separated by a narrow synaptic cleft of less than 50 nm. At these synapses, the release of neurotransmitter is due to the docking and fusion of vesicles to the presynaptic membrane through a mechanism triggered by Ca^2+^ influx into the presynaptic terminal, activation of the Ca^2+^ sensor synaptotagmin 1 (Syt1) (1) and recruitment of Soluble N-ethylmaleimide sensitive factor attachment protein receptor (SNARE) complex proteins (for review, see (2)). Docking and fusion of synaptic vesicles occurs at domains called active zones (AZ), that contain scaffolding proteins including Bassoon, RIM1/2 and ELKS (3). The identity of the proteins that are necessary and sufficient to establish the tight pre- and postsynaptic configuration of synapses is still unclear. However, synapse formation is regulated by trans-synaptic proteins including neurexins (Nrxn) and neuroligins (NL). The NL family comprises postsynaptic adhesion molecules that form trans-synaptic contacts with presynaptic neurexins (4–9)

The axonal connectivity of the dopamine (DA) system is particularly intriguing due to the hyperdense nature of its axonal arbor, containing a very large number of release sites, and to the ability of some of these neurons to release glutamate and GABA at a small subset of these (10–16). Strikingly, ultrastructural examination of the axon terminals established by monoamine and cholinergic neurons in the CNS of rodents, primates and humans failed to identify a tight pre- and postsynaptic coupling at most release sites, giving rise to the concept of “diffuse” or “volume” transmission, whereby neurotransmitter release from non-synaptic axon terminals leads to activation of metabotropic receptors on target cells located at a distance, within a sphere of a few tens of microns (17–24).

The molecular mechanisms determining the ability of DA neurons to establish synaptic and non-synaptic terminals are presently unknown. Midbrain DA neurons from the ventral tegmental area (VTA) project to the ventral striatum (VS) and to the prefrontal cortex (PFC), whereas DA neurons from substantia nigra (SNc) project densely to the dorsal striatum (DS) (12,25,26). The striatum is also densely innervated by glutamate neurons from the cortex that are thought to establish purely synaptic contacts. Interestingly, formation of excitatory glutamate synapses are mainly regulated by the presynaptic protein Nrxn-1 and the postsynaptic protein NL-1 (8, 9, 27–30), raising the hypothesis that the small subset of synaptic terminals established by DA neurons also depends on this pair of trans-synaptic proteins.

Here we developed an *in vitro* primary culture system to test the hypothesis that DA neurons are characterized by an intrinsic and distinctive propensity to establish an axonal arbor with limited synaptic connectivity. With this system, comparing DA neurons with cortical and striatal neurons, we confirm the unique ability of DA neurons to develop a majority (~80%) of non-synaptic terminals. Strikingly, non-synaptic axonal varicosities were found to be distinctive from the small subset of synaptic terminals in that most of them lack the active zone scaffolding proteins bassoon and RIM. The majority (~80%) of both synaptic and non-synaptic varicosities were found to be functional. Finally, using an artificial synapse formation assay, we found that NL-1 and Nrxn-1α play a critical role in the formation of synapses by DA neurons.

## MATERIALS AND METHODS

### Animals

All procedures involving animals and their care were conducted in accordance with the Guide to care and use of Experimental Animals of the Canadian Council on Animal Care. The experimental protocols were approved by the animal ethics committees of the Université de Montréal. Housing was at a constant temperature (21°C) and humidity (60%), under a fixed 12-h light/dark cycle with food water available ad libitum.

### Primary neuronal co-cultures

Primary DA neurons of VTA and SNc were prepared from transgenic mice expressing the Green Fluorescent Protein (GFP) gene in catecholamine neurons under control of the tyrosine hydroxylase (TH) promoter [(TH-GFP mice; (31)] and placed in co-culture with VS or DS neurons, respectively, prepared from C57BL/6J mice or from transgenic mice expressing the GFP protein fused to the DA D2 receptor(D2R) [D2R knock in mouse, GFP-D2 (32)]. Depending on the experiment, mesencephalic DA neurons from TH-GFP mice were selected by fluorescence-activated cell sorting (FACS) as previously described (33). Primary cultures of olfactory bulb (OB) DA neurons from TH-GFP mice were also prepared. As a point of comparison with DA neurons, we also prepared primary cultures of cortical glutamatergic neurons from C57BL/6J mice and dorsal striatal neurons from transgenic mice expressing GFP in neurons containing the DA D2 receptor (D2-GFP) (32, 34). For artificial synapse co-culture assays with HEK293T cells, primary DA neurons from VTA and SNc were prepared from transgenic mice conditionally expressing the red fluorescent protein tdTomato in DA neurons (DATcre-AI9 mice). For all experiments, postnatal day 0-3 (P0-P3) mice were cryoanesthetized, decapited and used for co-cultures and cultures according to a previously described protocol (35) (**Fig. S1**).

### Immunocytochemistry on cell cultures

Cultures were fixed at 14-DIV with 4% paraformaldehyde (PFA; in PBS, pH-7.4), permeabilized with 0,1% triton X-100 during 20-min, and nonspecific binding sites were blocked with 10% bovine serum albumin during 10-min. Primary antibodies were: chicken anti-GFP (1:2000, Aves Laboratories), rabbit anti-Syt1 (1:200, Synaptic Systems), rabbit anti-VMAT2 (1:4000, gift of Dr. Gary Miller, Colombia University), rat anti-DAT (1:5000; Chemicon) and guinea-pig anti-VGluT1 (1:4000, Abcam), mouse anti-HA (1:500; Roche). To improve the immunoreactivity of some synaptic markers, a set of cultures were fixed with 4% PFA together with 4% sucrose, in PBS without potassium (pH-7.4). Primary antibodies were, mouse anti-PSD95 (1:1000 Pierce antibody), mouse anti-gephyrin (1:1000, Synaptic Systems), guinea pig anti-bassoon (1:1000, Synaptic Systems), rabbit anti-RIM-1/2 (1:1000, Synaptic Systems). These were subsequently detected using Alexa Fluor-488-conjugated, Alexa Fluor-546-conjugated, Alexa Fluor-568-conjugated and Alexa Fluor-647-conjugated secondary antibodies (1:500, Invitrogen).

### Image acquisition with confocal microscopy

All *in vitro* fluorescence imaging quantification analyses were performed on images acquired using an Olympus Fluoview FV1000 point-scanning confocal microscope (Olympus, Tokyo, Japan). Images were scanned sequentially to prevent non-specific bleed-through signal using 488, 546 and 647-nm laser excitation and a 60X (NA 1:42) oil immersion objective.

### Image analysis

All image quantification was performed using ImageJ (National Institute of Health, NIH) software. A background correction was first applied at the same level for every image analyzed before any quantification. A macro developed in-house was used to perform all quantifications except the bassoon distribution analysis, performed with the 3D Roi-Manager plugin.

### FM1-43 loading and destaining

To quantify if DAergic terminals were capable of activity-dependent neurotransmitter release and vesicular cycling, we used the FM1-43 dye (ThermoFisher Scientific). FM1-43 release in response to KCl depolarization (40-mM) was monitored by time-lapse imaging using an epifluorescence microscope (Nikon TE2000) equipped with a Hamamatsu flash camera. FM1-43 labeled terminals were imaged using blue light 488-nm excitation and a 520-nm bandpass emission filter. High KCl-induced destaining of FM1-43 was performed by switching the perfusion solution (containing in mM: NaCl, 140; KCl, 5; CaCl2, 2; MgCl2, 2; HEPES, 10; glucose, 10; sucrose, 6) to a saline solution containing 40-mM KCl for 2-min. Images were acquired every 15s, with a 450-ms exposure time using a 100x 1.30 numeric aperture (NA) oil objective (Olympus, Tokyo, Japan).

### FM1-43 image analysis and quantification of KCl-induced FM1-43 destaining

After each recording, coverslips were fixed with 4% PFA and stained for GFP, Syt1 and VMAT2 to confirm that the experiment was performed exclusively on axonal varicosities established by DA neurons. GFP terminals positive for Syt1 were considered as potential DA release sites and included in the analysis. GFP terminals negative for Syt1 were considered as potentially non-competent and excluded from the analysis. With the ImageJ software, a region of interest (ROI) was created for each GFP varicosity positive for Syt1 to quantify signal intensity during the time-lapse. The GFP signal was subtracted from each picture to keep only the FM1-43 signal. Axonal varicosities established by GFP-positive neurons were considered FM1-43 positive when their fluorescence intensity was 2-fold greater than the background signal. We measured FM1-43 fluorescence level before and after KCl depolarization for each GFP varicosity positive for Syt1. Only puncta destaining by more than 20% were considered as active varicosities. For all data presented, at least 6 coverslips were analyzed from VTA DA neurons and 6 coverslips from SNc DA neurons. Each coverslip was only used for one experiment.

### Scanning electron microscopy

#### Titanium disc preparation

Commercially pure grade II titanium discs (Firmetal Co., Ltd., Shanghai, China) of approximately 12-mm diameter and 2-mm thickness were polished, sonicated in distilled water and air-dried (36). Before seeding the cells, discs were sterilized using 70% ethanol and UV light.

#### Morphological analysis

Neurons were cultured on titanium discs at a cell density of 60 000 cells/ml. After a period of 14-DIV, cells were fixed for 30-min at 4°C in 0.5% glutaraldehyde (EMS, Hatfield, USA). Samples were then rinsed three times with 0.1M cacodylate buffer (pH-7.4), and post-fixed for 1-h in 1% osmium tetroxide at 4°C. The cells were then dehydrated through a graded ethanol series (30%, 50%, 70%, 90%, 95%, and two times 100%; 15-min each) followed by critical point drying using a Leica EM CPD300 (Leica EM, Morrisville, USA). Characterization of cell morphology was carried out in a JEOL JSM-7400F (JEOL Ltd, Tokyo, Japan) field emission scanning electron microscope (FE-SEM) operated at 2-kV.

### Transmission electron microscopy (TEM)

#### Culture preparation

Cells were cultured in a Lab-Tek II permanox chamber slide system (ThermoFisher) at 240 000 cells/ml (120 000 cells/ml from VTA or SNc with 120 000 cells/ml from ventral striatum or dorsal striatum). Cells were fixed at 14-DIV for 1-h in 1% glutaraldehyde (EMS, Hatfield, USA) diluted in 0.1M cacodylate buffer. To identify specifically the ultrastructure of DA varicosities we performed a 3,3′-diaminobenzidine tetrahydrochloride (DAB) immunostaining against GFP protein. Sections were first rinsed in 0.1-M cacodylate, then in 0.1-M cacodylate + 30% H2O2 for 10-min. Neurons were then rinsed three times in 0.1-M cacodylate. Sections were then incubated in rabbit GFP antibody (1:2000; Abcam) for 24-h at RT. After rinses in 0.1-M cacodylate (3X for 10-min), neurons were incubated for 3-h at room RT in biotin-streptavidin conjugated AffiniPure goat anti-rabbit IgG (1:200; Jackson Immunoresearch), washed three times with 0.1 M cacodylate and then incubated for 3-h at RT in streptavidin HRP conjugate (1:200; GE Healthcare). Sections were visualized after 1-min and 30-sec DAB (Sigma-Aldrich)/glucose oxidase reaction. Next, cells were incubated in osmium solution with potassium ferrocyanide for 1-h (2% each) and rinsed with distilled water (3X for 10-min). Cells were dehydrated through an ethanol series (two times 30%, 50%, 70%, 90%, 95%, and two times 100%; 10-min each) and incubated in a solution containing 50% of Durcupan^®^ ACM Resin (Component A, 48.78%; Component B, 48.78%; Component C, 1.46% and Component D, 0.98%) and 50% of ethanol 100% for 1-h. Cells were then incubated in pure solution of Durcupan^®^ ACM Resin overnight at room temperature. Then, 12-h later, cells were incubated at 60°C for 48-h allowing polymerization of the resin.

#### Ultrastructural observations

Samples were unmolded, and the resin side was trimmed to obtain a flat and thin layer of resin parallel to *in vitro* neuron and astrocyte layers. Samples were glued on the tip of a resin block, with the astrocyte layer left accessible. After being glued, samples were cut at 80nm with an ultramicrotome (model EM UC7, Leica). Ultrathin sections were collected on bare 150 mesh per inch copper grids, stained with lead citrate and examined with a transmission electron microscope (Tecnai 12; Philips Electronic, 100 kV) and an integrated digital camera (XR41, Advanced Microscopy Techniques).

### Stochastic Optical Reconstruction Microscopy (dSTORM)

Cultures were fixed for 30-min in 4% PFA and washed with 0,1M PBS (3 × 5min). Following permeabilization with 0,1% triton for 20-min, primary antibodies for GFP and PSD95 were applied for 1-h. dSTORM imaging was performed, essentially as described in (37) using a Nikon N-STORM system. Briefly, imaging was carried out in a buffer containing 10% (w/v) glucose, 1% (v/v) beta-mercaptoethanol, 50 mM Tris-HCl (pH 8), 10 mM NaCl, 34 μg/ml catalase, 28 μg/ml glucose oxidase. Two-color dSTORM images of synaptic and non-synaptic DAergic varicosities were acquired by labeling cultured mesencephalic neurons derived from TH-GFP mice. GFP was labeled with CF568 and PSD-95 with Alexa647. During image acquisition, the 561 nm and 647 nm lasers were held constant at 0.6- and 1.1-kW cm^−2^, respectively, while a 405 nm laser was gradually increased to <0.1 kW cm^−2^ to switch on only a fraction of the fluorophores to give an optically resolvable set of active fluorophores. For each image, 20000 cycles of one frame of 561 nm laser activation followed by one frame of 647 nm laser were acquired at a rate of 16 Hz per cycle. Localizations were fit with fit3Dcspline and drift was corrected for with redundant cross correlation as previously described (38, 39). Localization and visualization of single particle (ViSP) were done as previously published (40).

### Construction of expression vectors

The NL expression vector contained cDNA encoding the mature form of full length of Human NL-1 with splice inserts A and B with an added extracellularly HA-tagged, under control of a cytomegalovirus enhancer fused to the chicken beta-actin (CAG) promoter in a pcDNA3 vector. The extracellular HA-tagged CD4 control construct was expressed from a CMV promoter, also in a pcDNA3 vector. The plasmid encoding ECFP-tagged ectodomains of mouse Nrxn 1α (splice variant SS4-) under control of a CMV promoter in a pECFP-N1 vector was obtained from Dr. Ann-Marie Craig (University of British Colombia). To overexpress Nrxn 1α^SS4-^ specifically in neurons, the CMV promoter was excised and replaced by the human synapsin (hSYN) promoter. The ECFP tag was also replaced by DsRED2. The sequence of all plasmids was verified by Sanger sequencing. For overexpression of Nrxn 1α^SS4-^, a plasmid containing the full sequence of Nrxn 1α^SS4-^ was packaged into a lentiviral vector under control of the CMV promoter (Viral core, Université Laval, Qc, Canada). For these experiments, primary DA neuron co-cultures were infected 3 days after plating. Cells were then fixed and immunolabelled at 14DIV.

### Artificial synapse formation assay

Artificial synapse formation assays were performed using HEK293T cell line as described previously (41). HEK293T cells were maintained in DMEM supplemented with FBS and penicillin/streptomycin and transfected using TransIT-LT1 (Mirus Bio. LLC) with the plasmid containing the sequence of NL-1 or CD4 and incubated overnight at 37°C. HEK293T cells were trypsinized (0,05%), washed and then co-cultured (30 000 cells/mL) with VTA or SNc DA neurons (120 000 cells/mL) from P0-P3 DATcre-AI9 pups at 9DIV. HEK293T cells were also co-cultured (30 000 cells/mL) with cortical neurons (120 000 cells/mL) as a positive control for the artificial synapse formation assay. Co-cultures were fixed 24h later (10DIV) with 4% PFA for 30 min, followed by permeabilization with 0.1% triton for 20 min and incubation with blocking solution (5% BSA) for 10 min. Immunolabelling was performed as described previously.

### Reverse Transcriptase-quantitative PCR

We used RT-qPCR to quantify the amount of mRNA encoding Nrxn 1α and Nrxn 1ß and to validate the overexpression of Nrxn 1α in primary neurons. Coverslips containing neurons were trypsinized (2.5%) at 14 DIV and then RNA extraction was performed with the RNAeasy Mini Kit (Quiagen) according the manufacturer’s instructions. The concentration and purity of the RNA from DA neurons were determined using a NanoDrop 1000 (Thermo Scientific, Waltham, MA USA). Total purified RNA (40 ng) was reverse-transcribed in a total of 20 μl including 1 μl of dNTP, 1 μl of random hexamer, 4 μl of 5X buffer 5X, 2 μl of dithiothreitol (DTT), 1 μl of RNAse-Out and 1 μl of the Moloney Murine Leukemia Virus reverse transcriptase enzyme (MML-V). Quantitative PCR was carried out in a total of 15μl consisting of 3μl cDNA, 7.5μl SYBER green PCR master mix (Quanta Biosciences), 10μM of each primer, completed up to 15μl with RNA-free water. qPCR was performed on a Light Cycler 96 machine (Roche) using the following procedure: 10 min at 95°C; 40 cycles of 30s at 95°C, 40s at 57°C and 40s at 72°C; 1 cycle of 15s at 95°C, 15s at 55°c and 15s at 95°C. Results were analysed with Light Cycler 96 software and Excel. The efficiency of the reaction (E=10^(−1/slope)^ −1) was calculated from the slope of the linear relationship between the log values of the RNA quantity and the cycle number (Ct) in a standard curve. Calculation of relative mRNA levels was performed by using the 2^(-ΔΔCt) formula (42), where the Ct value of the mRNA level for Nrxn1 was normalized to the Ct value of GAPDH in the same sample. Ct values used were the mean of duplicate repeats. Melt-curves of tissue homogenate indicated specific products after Nrxn 1α and Nrxn 1 ß qPCR mRNA amplification, attesting of the adequate quality of the primers chosen. Primers for Nrxn 1α were obtained from the Sudhöf Lab and primers for GAPDH were designed with the Primers 3 and Vector NTI software. Primers were synthetized by Alpha DNA (Montreal, QC). Primers for qPCR were as follows: Nrxn 1α: 5’ CAACACAAATCACTGCGGG 3’ and 5’ TTCAAGTCCACAGATGCCAG 3’; Nrxn 1ß: 3’ TTGCAATCTACAGGTCACCAG 5’ and 3’CCTGTCTGCTGTGTACTG5’; GAPDH 5’ GGAGGAAACCTGCCAAGTATGA 3’ and 5’ TGAAGTCGCAGGAGACAACC 3’. Primers were tested by comparing primers sequences to the nucleotide sequence database in GenBank using BLAST (www.ncbi.nlm.nih.gov/BLAST/).

### Statistics

Data were always obtained from three separate sets of experiments and presented as mean ± standard error of mean (SEM). The level of statistical significance was established at p < 0,05 in one-way ANOVAs and two-tailed t tests, performed with Prism 6 software (GraphPad, ^*^p < 0.05, ^**^p < 0.01, ^***^p < 0.001, ^****^p < 0.0001)

## RESULTS

### Dopaminergic neurons in vitro establish mostly non-synaptic terminals

Previous work using transmission electron microscopy (TEM) demonstrated that in the intact rodent brain, the axon terminals of DA neurons in the striatum are only rarely found in synaptic contact with target cells, the directly apposed cellular membrane being typically devoid of a post synaptic density (PSD) (43). Whether this represents an intrinsic property of mesencephalic DA neurons or requires complex signals received *in vivo* during development is unknown. To begin exploring this key question we first used scanning electron microscopy (SEM) to visualize the external morphology of axonal varicosities in co-cultures of VTA or SNc DA neurons, obtained from TH-GFP transgenic mice, growing with VS or DS neurons, respectively, as well as astrocytes. Neurons were grown on surface-optimized titanium disks and examined by SEM after 14 days (**Fig. 1A** and **1B**). We observed that in such cultures, some of the neurons, likely dopaminergic, established a complex array of oblong axonal varicosities in close contact with the underlying astrocytes, but without direct contact with dendrites or neuronal cell bodies (**Fig. 1D** and **1E**). These varicosities have an average length of 1.4 μm with an average inter-varicose interval of 2.6 μm, highlighting the high density of varicosities along the axon (**Fig. 1D**). We found that only a very small contingent of varicosities along such axons established synaptic-like contacts with neuronal dendrites (**Fig. 1F**). Globally in these co-cultures, approximately 15% of neurons developed axonal arbors with a majority of non-synaptic contacts, in line with the typical proportion of DA neurons in such cultures. To confirm that this apparently non-synaptic connectivity originated from DA neurons, we prepared a second set of cultures in which FACS-purified DA neurons were grown without striatal neurons. In such monocultures, we observed a large predominance of free axonal varicosities (**Fig. 1G** and **1H**), confirming that DA neurons have an intrinsic propensity to develop a large and complex axonal arbor, with most varicosities being non-synaptic. As a point of comparison, we also examined striatal cultures, containing more than 95% GABAergic medium spiny neurons. In this system we found a large predominance of axonal varicosities establishing synaptic-like contacts, with most axons being non-varicose and containing only a small subset of free axonal varicosities (**Fig. 1I** and **1J)**. Although accounting for less than 5% of striatal neurons, some of these free varicosities could also originate from the axons of striatal cholinergic neurons, previously reported to also establish mostly non-synaptic terminals in the striatum (21).

**Figure 1.**
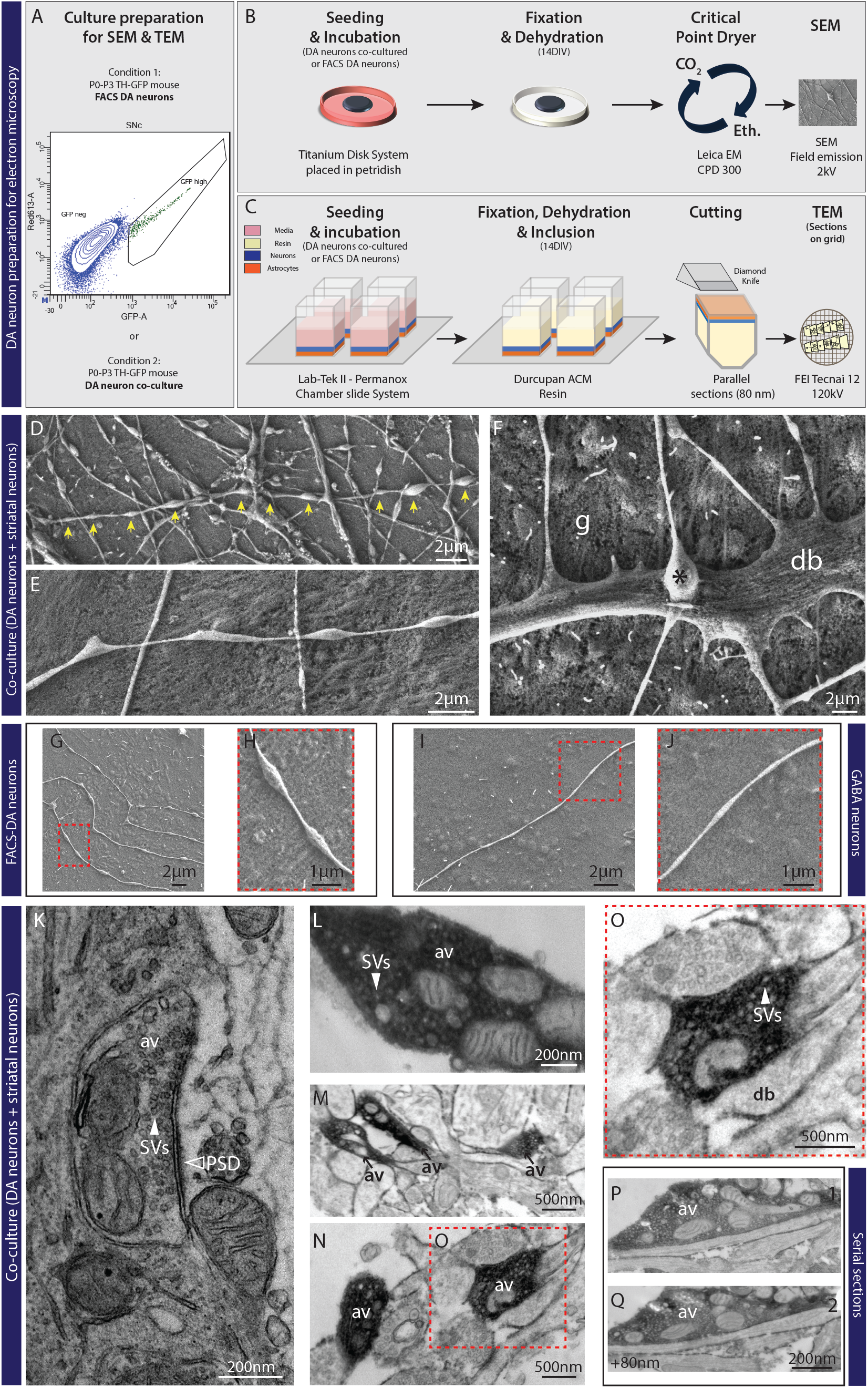
Ultrastructural evidence for functional non-synaptic terminals established by DA neurons. **A to C**– Schematic representation for SEM and TEM experiments (see materials and methods for details). Graph-plot for FACS experiments showing the cells selected based on a high level of GFP signal. **D** and **E**– Electron micrographs from SEM showing free varicosities (yellow arrows) established by VTA neurons co-cultured with ventral striatal neurons. **F**- Electron micrograph from SEM showing a direct apposition between a terminal (*) and a dendrite (d), in close interaction with glial cells (g). **G** and **H**- Electron micrographs from SEM showing non-synaptic DA varicosities from FACS-purified DA neurons. **I** and **J**- SEM image from an axonal field of a GABA neuron showing the absence of free varicosities compared to DA neurons. **K**- Electron micrograph from TEM showing a synaptic terminal established by a non-DA neuron, associated with a PSD domain (black arrow) and filled with synaptic vesicles (SVs). **L**- Electron micrograph from TEM showing a DA immunoreactive varicosity containing a substantial pool of vesicles (see white arrow) and a cluster of 4 mitochondria. **M** and **N**- Electron micrographs from TEM showing GFP immunoreactive non-synaptic DAergic varicosities (AV) established by VTA DA neurons co-cultured with ventral striatal neurons. **O**- Electron micrograph from TEM. **P** and **Q**- Serie of 2 consecutive ultra-thin sections across a GFP immunoreactive non-synaptic DAergic varicosity established by a VTA-DA neuron co-cultured with ventral striatal neurons.

We then used TEM (**Fig. 1C**) to visualize the internal organization of axonal varicosities in co-cultures of VTA or SNc DA neurons, growing with VS or DS neurons, respectively, on astrocytes. Using 3,3′-diaminobenzidine tetrahydrochloride (DAB), we performed immunostaining against GFP to distinguish DA and non-DA terminals. In our *in vitro* system, we easily found DAB negative synaptic terminals, identified by the presence of a post-synaptic domain as illustrated in **Fig. 1K**. We then identified multiple sets of DAergic terminals, all filled with large pools of synaptic vesicles, arguing that they possess the capacity to package and release neurotransmitters (**Fig. 1L, 1M** and **1N**). However, none were found in close proximity to a post-synaptic domain, revealing the asynaptic nature of DAergic terminals (**Fig. 1O**). Due to the possibility that some of these terminals may show a post-synaptic domain in another plane of the varicosity, we searched for the same axon varicosity on different ultrathin sections. Even in such case, no post-synaptic domains were observed in close proximity to DAergic terminals (**Fig. 1P** and **1Q**).

### Non-synaptic dopaminergic axonal varicosities express distinct sets of presynaptic markers and appear to actively avoid target cells

Although SEM and TEM analysis revealed the non-synaptic nature of DAergic axonal varicosities, it was unclear if all such morphologically identified varicosities represented actual axon terminals endowed with the molecular machinery required for neurotransmitter release. We next used immunocytochemistry coupled to confocal microscopy to examine the axonal varicosities established by DA neurons from TH-GFP mice. Double labelling for GFP, to visualize all varicosities and the ubiquitous Ca^2+^ sensor for exocytosis Syt1, we found that 89.0 ± 2.5% of varicosities established by SNc DA neurons were positive for Syt1 (**Fig. 2A** and **2F**). This proportion was slightly smaller for VTA DA neurons (69.3 ± 2.8%) (**Fig. 2F**). To gain further insight into the neurochemical identity of these axon terminals, we next examined the presence of the vesicular monoamine transporter VMAT2, necessary for the vesicular packaging and release of DA and the type 2 vesicular glutamate transporter VGluT2, necessary for glutamate release by DA neurons (14, 44–46). Double-labeling DA neuron co-cultures for GFP and VMAT2 revealed that 50.1 ± 6.1% and 53.8 ± 5.4% of SNc and VTA DA neuron terminals, respectively, contained VMAT2 (**Fig. 2B** and **2G**). Similar double-labeling experiments evaluating VGluT2 expression revealed a small proportion of GFP-positive glutamatergic terminals established by DA neurons (**Fig. S2**). These results indicate that most GFP-positive axonal varicosities are likely to represent sites allowing the release of DA, as well as other neurotransmitters by DA neurons.

**Figure 2.**
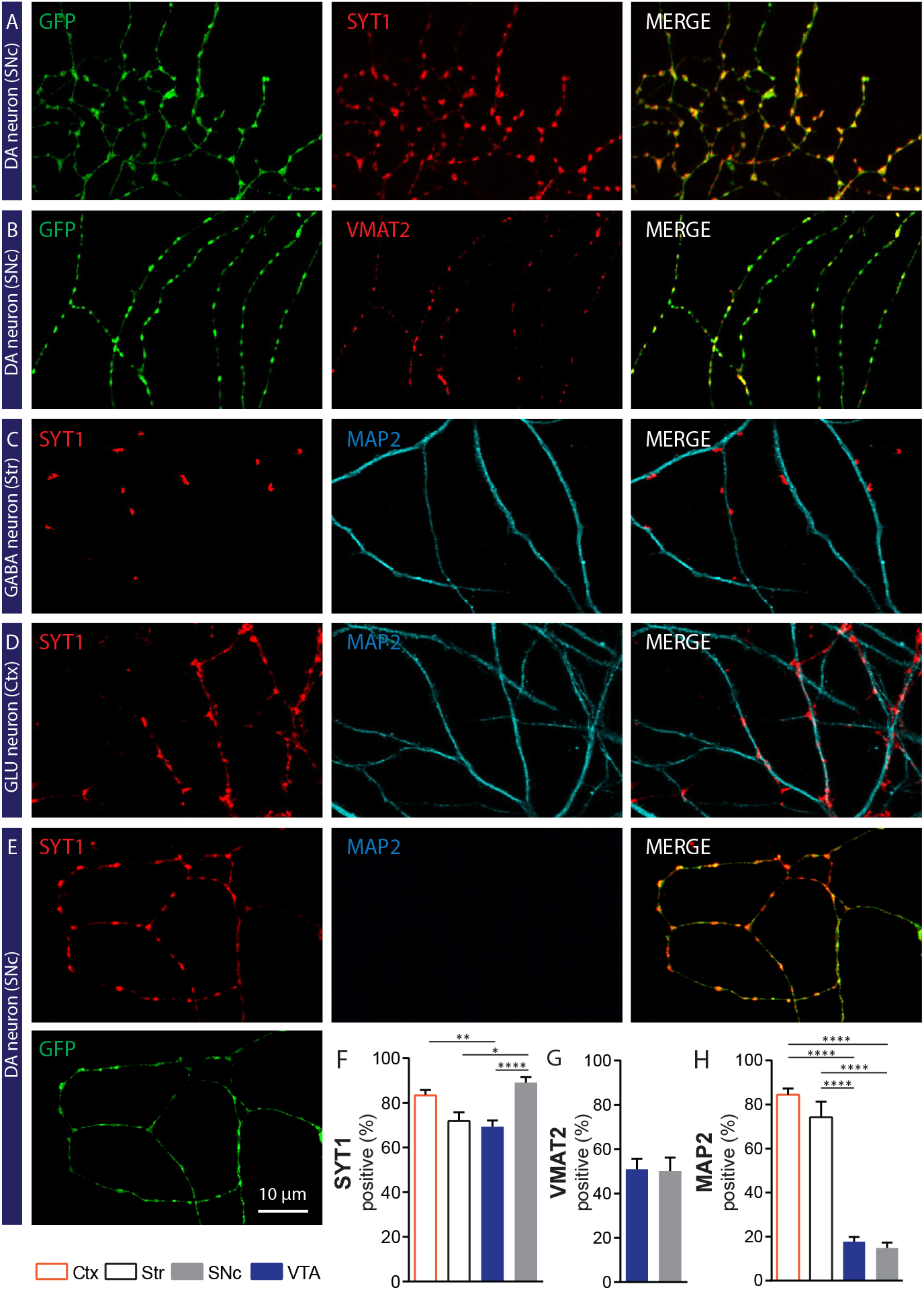
Dopaminergic neurons establish a majority of axon terminals that are not in contact with target cells. **A** and **B**- Photomicrographs illustrating a majority of GFP-positive axonal varicosities established by a SNc DA neuron, that contain Syt1 (Panel **A**) or VMAT2 (Panel **B**). **C**- Photomicrographs illustrating Syt1 positive terminals established by a GABAergic striatal neuron in contact with a MAP2-positive dendrite from a target cell. **D**- Photomicrographs illustrating Syt1 positive terminals established by a glutamatergic (GLU) neuron in a cortex-striatum co-culture, in contact with a MAP2-positive dendritic process from a target cell. **E-** Photomicrographs illustrating GFP terminals positive for Syt1 established by a SNc DAergic neuron in co-culture with striatal neurons, in a field without any target cell. **F**- **G**- Bar graphs representing the quantification of the proportion (%) of GFP positive varicosities that co-express Syt1 (**F**) or VMAT2 (**G**), for both VTA and SNc DA neurons. Bar graphs representing the mean +/− SEM, one-way ANOVA, Tukey’s multiple comparison test, ^*\*\*\*\**^*p<0.0001*. For Syt1 analysis: Cortex n=36, Striatum n=25, VTA n=59, SNc n=46; from 3 different cultures. For VMAT2 analysis: VTA n=29, SNc n=28; from 2 different cultures. The number of observations represents the number of fields from individual neurons examined. **H**- Bar graphs representing the proportion of Syt1 varicosities established by VTA or SNc DAergic neurons that are in close proximity to MAP2-positive dendrites, compared to glutamatergic and GABAergic neurons from cortex and striatum, respectively. Bar graphs represent the mean +/− SEM, one-way ANOVA, Tukey’s multiple comparison test, ^*\*\*\*\**^*p<0.0001*. Cortex n=20; Striatum n=19; VTA n=39; SNc n=25; from 2 different cultures. The number of observations represents the number of fields from individual neurons examined.

As our results to this point suggested that most DAergic terminals were non-synaptic in nature and thus tended to spontaneously avoid interacting with the otherwise numerous striatal medium spiny neurons, we next aimed to examine more directly the interaction of DAergic terminals with the somatodendritic domain of medium spiny neurons, visualized here using a MAP2 antibody. Using triple-labeling for GFP, Syt1 and MAP2, we compared random fields of SNc or VTA DAergic axonal domains with axonal fields acquired from striatal and cortical cultures, thus providing a comparison with neuronal populations known to be essentially completely synaptic in their connectivity (**Fig. 2C to 2E**). We calculated the proportion of GFP/Syt1 double-positive varicosities that were also in direct contact with MAP2-positive elements, thus providing an index of the propensity of terminals to be in close contact with the dendritic or somatic membrane of target cells. Strikingly, we found that only 14.9 ± 2.4% of axon terminals established by SNc DA neurons were in contact with target cells (**Fig. 2C** to **2E**). This proportion was not significantly different for VTA DA neurons (17.8 ± 2.1 %) (**Fig. 2H**). In marked contrast, 84.5 ± 2.7% of terminals in cortical cultures (**Fig. 2D** and **2H**) and 74.2 ± 7.1% of terminals in striatal cultures (**Fig. 2C** and **2H**) were found to be in close proximity to target cells. These findings bring to light the very different mode of connectivity of DA neurons and their main target cells compared to that of classical glutamatergic and GABAergic neurons.

### Hyperdense nature of the axonal domain of DA neurons compared to glutamate and GABA neurons

Another intriguing aspect of the connectivity of DA neurons is the very dense, highly complex and branched nature of their axonal compartment (12, 15, 16, 47). As we examined the distribution of presynaptic markers along the axonal domain of DA neurons, we observed that the density of axonal varicosities along DAergic axons appeared to be higher compared to other types of neurons. We therefore quantified the distribution of Syt1-positive axonal varicosities along the axonal domain of VTA and SNc DAergic neurons and compared this to striatal GABA and cortical glutamate neurons (**Fig. 3A to 3C**). We found that the inter-varicosity distance was 9.0 ± 1.8μm and 10.9 ± 2.2μm for SNc and VTA DA neurons, respectively. This value was two-fold larger for GABA and glutamate neurons, for which the inter-varicosity distance was 22.1 ± 3.6μm and 27.6 ± 2.9μm, respectively (**Fig. 3D**). This observation further highlights the fundamentally different nature of dopaminergic axonal arbors compared to those of GABA and glutamate neurons.

**Figure 3.**
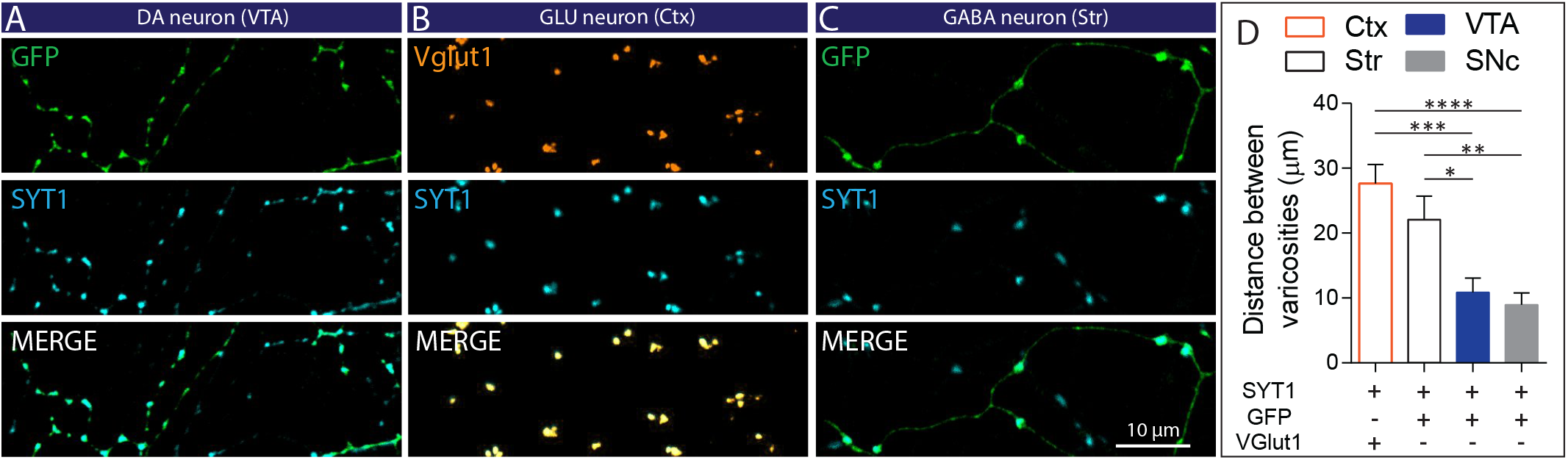
Dopaminergic neurons establish a high density of axonal varicosities along their axonal domain. **A**- Photomicrographs illustrating the distribution of GFP/Syt1 positive axonal varicosities along the axonal domain of a VTA DAergic neuron. **B**-Photomicrographs illustrating the distribution of VGluT1/Syt1 positive varicosities along the axonal domain of a glutamatergic (GLU) neuron from cortex. **C**- Photomicrographs illustrating the distribution of GFP/Syt1 positives axonal varicosities along the axonal domain of a GABAergic D2-GFP striatal neuron. **D**- Bar graphs representing the distance (μm) between Syt1 positive varicosities along the axonal domain of VTA and SNc DAergic neurons compared to glutamatergic and GABAergic neurons from cortex and striatum, respectively. Bar graphs present the mean +/− SEM, one-way ANOVA, Tukey’s multiple comparison test, ^*\**^*p<0.05;* ^*\*\**^*p<0.01;* ^*\*\*\**^*p<0.001;* ^*\*\*\*\**^*p<0.0001.* Cortex n=12; Striatum n=12; VTA n=12 SNc n=12; from 3 different cultures. The number of observations represents the number of fields from individual neurons examined.

### Limited expression of the active zone scaffolding proteins bassoon and RIM

Because axon terminals established by DA neurons clearly differed in their mode of connectivity compared to GABA and glutamate neurons, we sought to further examine some key active zone components known to be important for the neurotransmitter release machinery of neurons. At classical fast synapses, active zones are typically characterized by the expression of specific scaffolding proteins such as bassoon and RIM1/2 (48). Interestingly, recent work showed that only ~30% of DAergic terminals *in vivo* express the active zone proteins bassoon and RIM (49). However, it has not been determined whether synaptic and non-synaptic DA terminals differentially contain such scaffolding proteins. We hypothesized that synaptic DA terminals might be more likely to express active zone scaffolding proteins compared to the non-synaptic DA terminals because of the necessity to maintain a stable pre- and postsynaptic complex. Confirming previous results, we found that only 36.0 ± 5.5% and 32.2 ± 3.8% of SNc or VTA DA terminals were positive for bassoon (**Fig. 4A, 4D**). A striking observation was that the majority of bassoon positive DAergic terminals were either in direct contact or in close proximity to a target cell. In axonal fields far removed from other neurons, axonal varicosities were typically completely devoid of bassoon (**Fig. 4A**, upper panels, **4F**). In axons closer to striatal neurons (**Fig. 4A**, lower panels), approximately one third of varicosities in contact with MAP2-positive somatodendritic domains were bassoon positive (27.3 ± 4.5% and 28.9 ± 3.0% for SNc and VTA DA neurons, respectively; **Fig. 4E**). In these fields, bassoon was found in DA varicosities that were located between 0.5μm to 40μm from MAP2-positive dendrites, as illustrated schematically (**Fig. 4G** and **Fig. S3**).

**Figure 4.**
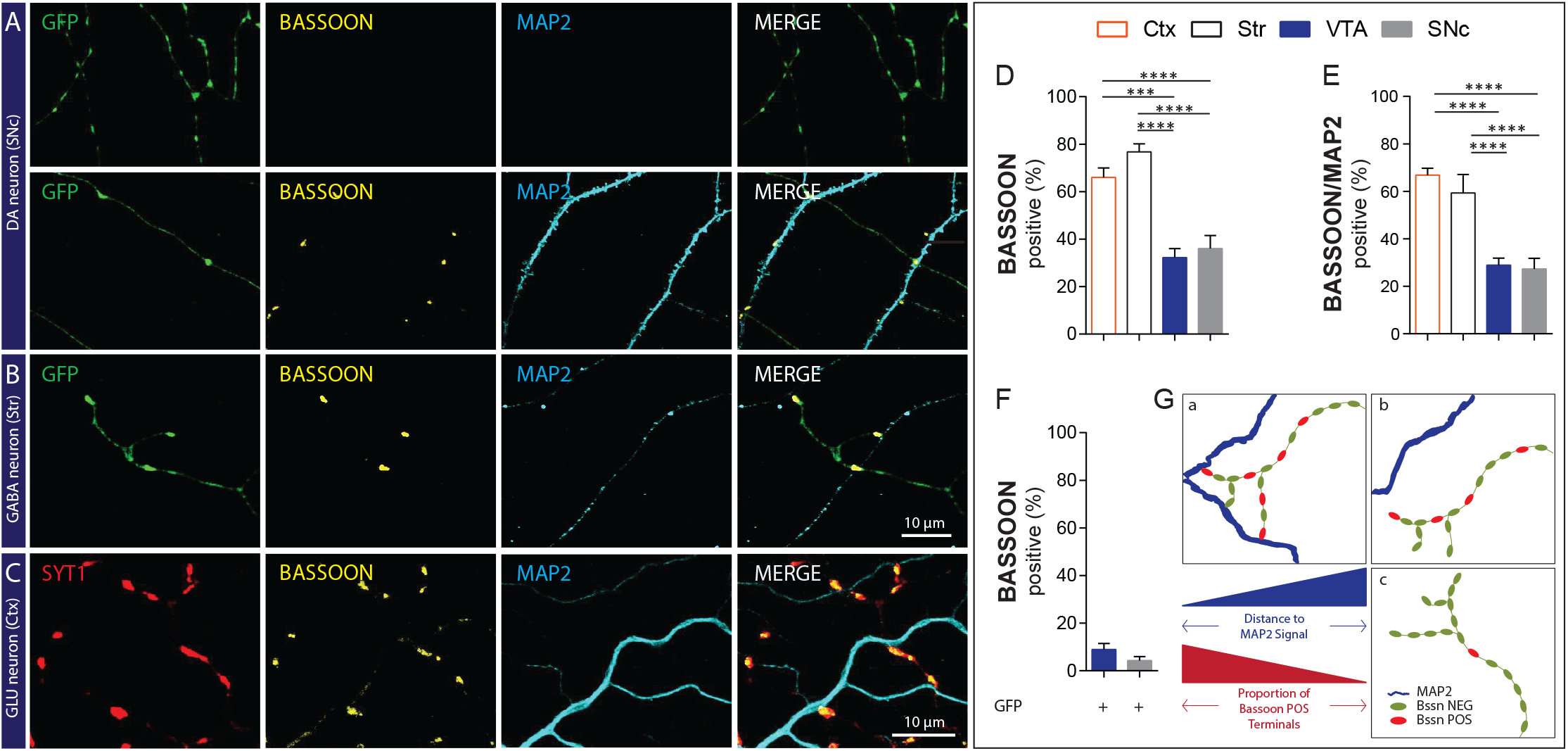
The active zone protein bassoon is enriched in DA terminals that are in close proximity to dendrites. **A-** The top set of photomicrographs illustrates that GFP positive axonal varicosities established by a SNc DA neuron are negative for bassoon and distant from any MAP2 positive dendrites. The bottom set of photomicrographs illustrates that a larger subset of GFP positive axonal varicosities established by a SNc DA neuron is positive for bassoon when localized close to MAP2 positive dendrites. **B**- Photomicrographs illustrating that most GFP positive axonal varicosities established by a striatal GABAergic neuron co-express bassoon and are localized close to MAP2 positive dendrites. **C**- Photomicrographs illustrating that most Syt1 positive axonal varicosities established by a cortical glutamatergic neuron co-expresses bassoon and are localized in proximity to MAP2 positive dendrites. **D**- Bar graphs representing the proportion of GFP positive axonal varicosities established by VTA and SNc DAergic neurons that co-express bassoon, compared to Syt1 positive axonal terminals from cortical glutamatergic neurons and striatal GABAergic neurons. **E**- Bar graphs representing the proportion of axonal varicosities established by VTA and SNc DAergic neurons that co-express bassoon and are in contact with MAP2 positive dendrites, compared to glutamatergic and GABAergic neurons from cortex and striatum, respectively. **F-** Bar graphs representing the proportion of axonal varicosities established by VTA and SNc DAergic neurons that co-express bassoon in a context of absence of a target cell in the field. **G**- Schematic representation of bassoon-positive DA terminals based on the localization of a target cell, represented by the MAP2 signal. Bar graphs represent the mean + SEM, one-way ANOVA, Tukey’s multiple comparison test, ^*\*\*\**^*p<0.001;* ^*\*\*\*\**^*p<0.0001*. Cortex n=40; Striatum n=16; VTA n=17; SNc n=22 from 2 different cultures. The number of observations represents the number of fields from individual neurons examined.

In striking contrast, most terminals established by cortical glutamate neurons were positive for bassoon (66.0 ± 3.9%) (**Fig. 4B, 4D**), while for striatal GABA neurons, this was 76.8 ± 3.4% (**Fig. 4C, 4D**). Furthermore, 66.8 ± 2.9 % of bassoon-positive glutamatergic terminals and 59.3 ± 7.8 % of striatal GABAergic terminals were in contact with MAP2-positive structures (**Fig. 4E**).

We found RIM1/2 to also be sparsely expressed by DAergic terminals, unrelated to the presence of target cells. While in some fields, good colocalization of RIM1/2 and bassoon was observed (**Fig. S4A**), in others, they were mostly segregated (**Fig.S4B** and **S4C**). Globally, 31.7 ± 5.1% and 38.9 ± 5.1% of SNc and VTA DAergic terminals contained RIM1/2 (**Fig. S4D**). The proportion of bassoon-positive terminals expressing RIM1/2 was 33.2 ± 5.4% and 29.6 ± 5.9%, respectively for SNc and VTA DA neurons (**Fig. S4E**). Conversely, 37.6 ± 6.3% and 25.7 ± 5.3% of bassoon positive SNc and VTA DAergic terminals contained RIM1/2 (**Fig. S4F**), highlighting their mostly segregated distribution.

### Synaptic contacts established by DA neurons are neurochemically diverse

Our results thus far highlight the surprising propensity of DAergic neurons to establish a majority of neurotransmitter release sites that not in direct contact with target cell dendrites are thus non-synaptic. This observation contrasts with the opposite propensity of cortical glutamate or striatal GABA neurons. The small subset of synaptic-like release sites established by DA neurons could represent sites implicated in glutamate and GABA release because previous work has described that glutamate and GABA can act as additional co-transmitters in DA neurons (11, 14, 44, 46, 50–52). To gain further insight into the synaptic-like terminals detected in our model, we next examined the relationship between DAergic terminals and postsynaptic organizers associated with glutamatergic (PSD95) and GABAergic (gephyrin) synapses (53, 54). We found that a small subset of Syt1-positive terminals established by SNc DA neurons was in close apposition to PSD95 (12.6 ± 3.3%) or gephyrin (5.2 ± 1.8%). A similar low proportion of VTA DA neuron terminals were found in close proximity to PSD95 (8.0 ± 2.3%) or gephyrin (2.9 ± 0.8%) (**Fig. 5A, 5E** and **5F**). The difference between SN and VTA was not significant. To visualize at higher resolution the molecular architecture of synaptic and non-synaptic terminals established by primary DA neurons, we used direct stochastic optical reconstruction microscopy (dSTORM). For the synaptic DA varicosities, the dSTORM images revealed a clear apposition between the DAergic varicosity, defined by GFP nanocluster signal, and the postsynaptic domain of a target cell, defined by PSD95 nanocluster signal (**Fig. 5H**). An example of non-synaptic DAergic terminal visualized by super-resolution imaging is also illustrated, demonstrating the lack of proximity to a PSD domain (**Fig. 5I**). In stark contrast, 83.6 ± 2.2% of VGluT1/Syt1 terminals established by cortical glutamate neurons colocalized with PSD-95 (**Fig. 5B** and **5E**). Similarly, the large majority (60.6 ± 3.7%) of terminals established by striatal GABAergic neurons were in close apposition to gephyrin postsynaptic clusters (**Fig. 5C** and **5F**). Because previous work has suggested that olfactory bulb (OB) DA neurons release GABA as a co-transmitter at many of their release sites, we also examined synaptic release sites from such neurons. Interestingly we found that 55.4 ± 5.2% of Syt1-positive DAergic terminals established by such neurons colocalized with gephyrin (**Fig. S5**).

**Figure 5.**
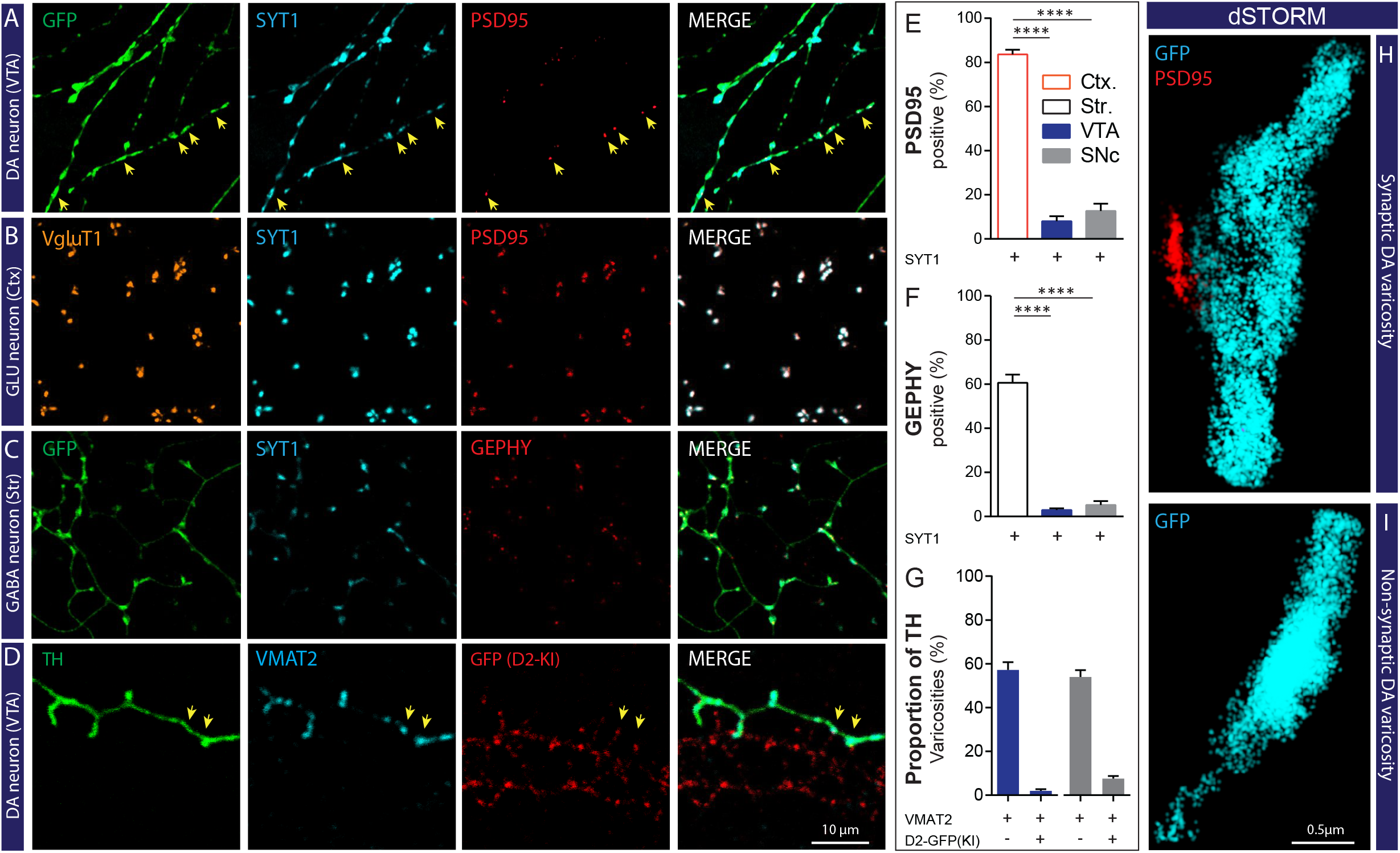
DA neurons develop only a small proportion of excitatory and inhibitory synapses in contrast to cortical and striatal neurons and a small subset of terminals in close proximity to D2R clusters. **A-** Photomicrographs illustrating GFP positives varicosities in the axonal arborization of a VTA DA neuron expressing Syt1 and colocalizing sparsely with the postsynaptic marker PSD95. **B-** Photomicrographs illustrating VGluT1 positives varicosities along the axonal arborization of a cortical neuron expressing Syt1 and colocalizing with PSD95. **C-** Photomicrographs illustrating GFP positives varicosities in the axonal arborization of a GABAergic neuron expressing Syt1 and colocalizing with the postsynaptic marker gephyrin. **D-** Photomicrographs illustrating TH positives varicosities in the axonal arborization of a SNc DA neuron expressing VMAT2 and colocalizing sparsely with the D2R (yellow arrow). **E-** Bar graph representing the proportion (%) of axonal varicosities established by VTA, SNc, cortical and striatal neurons that are positive for Syt1 and colocalizing with PSD95 at 14 DIV. **F**- Bar graph representing the proportion (%) of axonal varicosities established by VTA, SNc, cortical and striatal neurons that are positive for Syt1 and colocalizing with gephyrin at 14 DIV. **G**- Bar graphs representing the proportion (mean and SEM (%)) of TH-VMAT2 positive terminals colocalizing with D2R (GFP signal). **H**- **I** Representative dSTORM images showing a synaptic DA varicosity GFP-positive (cyan) (**H**) in close contact to a PSD95 domain (red) and a non-synaptic GFP-positive (cyan) DAergic varicosity (**I**) without any contact to a post-synaptic domain. The bar graphs represent the mean + SEM, one-way ANOVA, Tukey’s multiple comparison test, ^*\*\*\**^*p<0.001;* ^*\*\*\*\**^*p<0.0001.* PSD95/gephyrin experiments: Cortex n=25; Striatum n=25; VTA n=25 SNc n=25; from 3 different cultures. D2-KI experiments: VTA n=35 SNc n=33; from 4 different cultures. The number of observations represents the number of fields from individual neurons examined.

Although evidence for DA receptor clusters at synaptic sites in the brain is limited (23, 24), DA neurons could in principle also establish synaptic-like release sites that are in close apposition to postsynaptic DA receptors, independently from PSD-95 or gephyrin domains. To examine this, we took advantage of a knock-in mouse line expressing GFP-tagged D2 receptors and visualized release sites specialized for DA release using VMAT2 immunostaining. In co-cultures of WT DA neurons together with striatal neurons from GFP-D2 mice (**Fig. 5D)**, we found that 54.0 ± 3.1% of TH-positive terminals established by SNc DA neurons contained VMAT2 (**Fig. 5G**). Similarly, 57.2 ± 3.6% of TH-positive DA terminals established by VTA DA neurons were VMAT2 positive (**Fig. 5G)**. Arguing here again for a strong propensity of DAergic axons to establish release sites that fail to seek out target cells, we found that only 7.6 ± 1.2% of TH/VMAT2 terminals established by SNc DA neurons contacted D2 receptor clusters, while this proportion was 2.0 ± 0.8% for VTA DA neurons (**Fig. 5G**).

### FM1-43 imaging reveals a majority of active DA terminals

Considering the large heterogeneity of DAergic terminals identified in the present work, we wished to determine whether all or only a subset of morphologically defined axonal varicosities are functional and competent to release neurotransmitters by exocytosis. Here we used the activity-dependent uptake and release of the well-characterized endocytotic probe FM1-43, allowing us to examine activity-dependent vesicular cycling independently of the neurochemical phenotype of the varicosities. After a first round of FM1-43 uptake induced by high potassium induced membrane depolarization, a second round of membrane depolarization was used to identify terminals that showed activity-dependent release of FM1-43 (**Fig. 6A** and **6B**). A total of 1091 SNc and 543 VTA varicosities were examined. To isolate activity-dependent release from spontaneous release or probe bleaching, we used as a criterion a minimal release of 20% of the initially up taken FM1-43 in response to the second round of membrane depolarization (**Fig. 6C**). We found that 84.62 ± 4.60% of SNc and 77.58 ± 9.62% of VTA axonal varicosities, identified as potential release sites by the presence of Syt1, were active (**Fig. 6D**). Post-experimental immunostaining of the same fields that were imaged revealed that a majority of FM1-43 puncta along DAergic axons contained Syt1 or VMAT2 (**Fig. 6E**).

**Figure 6.**
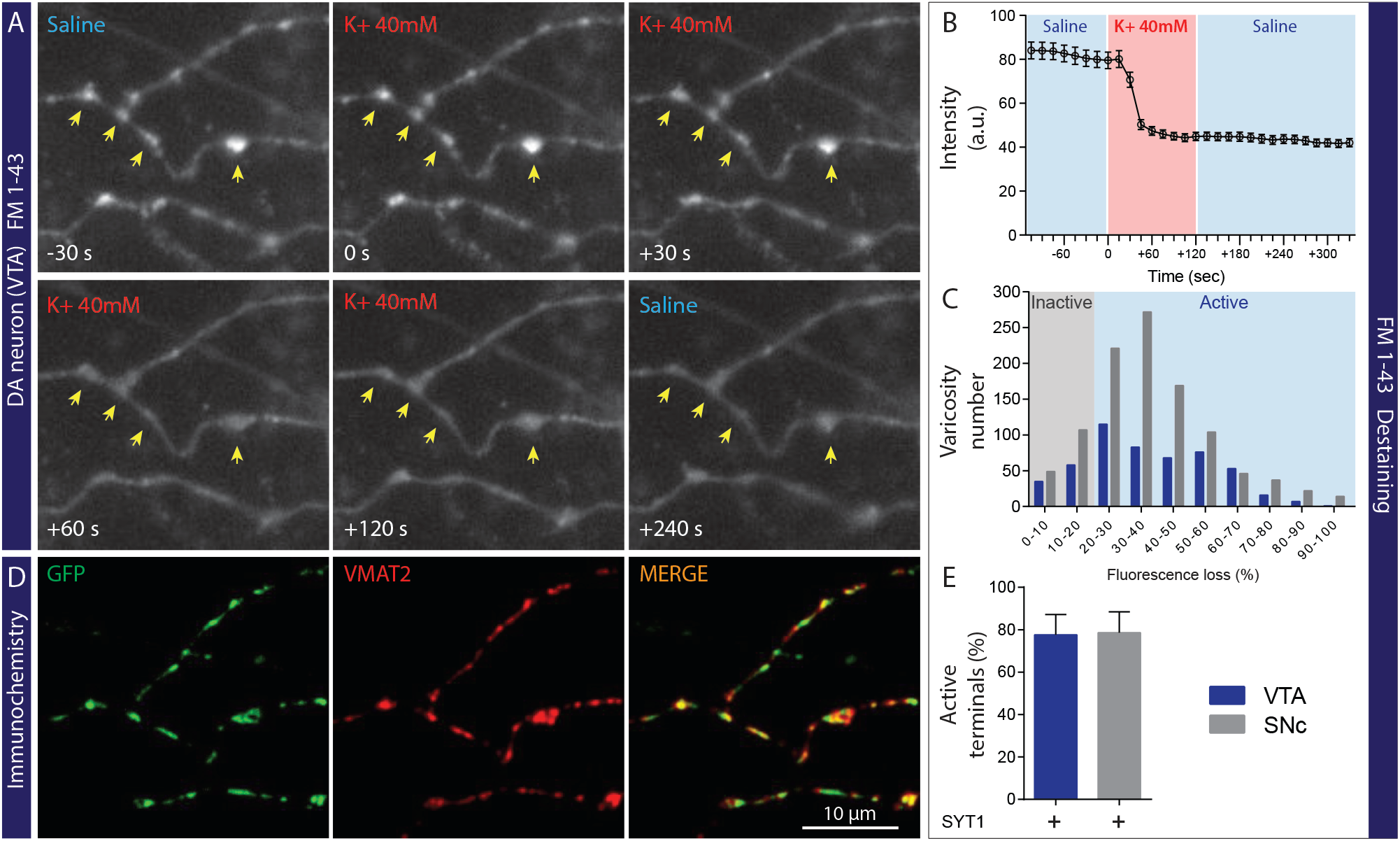
Most synaptic and non–synaptic dopaminergic terminals are active. **A**- Images of the axonal field from a VTA DA neuron labeled live with FM1-43. Representative images at the different indicated time points before, during and after KCl-induced membrane depolarization and FM1-43 destaining. Time 0 s represents the last image before KCl stimulation. **B**- Intensity (arbitrary units – a.u.) versus time plot for FM1-43 puncta that destained by at least 20% in response to 40mM KCl. **C**- Histogram showing the distribution of FM1-43 destaining levels in response to KCl depolarization (n=522 puncta from 5 different VTA DA neurons and n=986 puncta from 6 different SNc DA neurons). A loss of fluorescence < 20% was considered as an inactive varicosity. **D**- Images of the same axonal field illustrated in panel A after post-recording fixation and immunostaining for GFP and VMAT2. **E**- Bar graph representing the proportion (mean +/− SEM (%)) of active varicosities established by VTA and SNc DA neurons. Analysis including FM1-43 positive and FM1-43 negative varicosities (n=543 varicosities from 5 different VTA DA neurons; n=1091 varicosities from 6 different SNc DA neurons).

### NL-1 induces presynaptic differentiation of DA neuron terminals

Because a subset of the synapses established by DA neurons are glutamatergic and striatal neurons act as postsynaptic recipients for a massive number of glutamatergic synapses from the cortex, we hypothesized that the transsynaptic protein NL-1, specifically expressed at the postsynaptic compartment of excitatory synapses (28, 29, 55, 56), might play a role in regulating synapse formation by VTA and SNc DA neurons. We used an artificial synapse formation assay to test this hypothesis. Primary DA neurons derived from the VTA and SNc and expressing the red fluorescent protein tdTomato (RFP) were co-cultured with HEK293T cells expressing the major form of NL-1 (splice variant A^+^B^+^; NL-1^AB^) tagged with extracellular HA or a negative control membrane protein (CD4 with HA tag) and we examined whether DA axonal varicosities were recruited to establish synapse-like contacts (**Fig. 7A** and **7B**). We found that HEK293T cells expressing NL-1^AB^ were significantly more attractive for DA terminals, identified by the presence of RFP or VMAT2, compared to HEK293T cells expressing the control protein CD4 (**Fig. 7C**). Quantification of the total intensity of RFP and VMAT2 puncta on HEK293T cells expressing NL-1^AB^ showed a 5-fold and a 27-fold increase in signal, respectively, compared to the CD4 control group, representing background signal due to the random distribution of axons (**Fig. 7D** and **7E**). Cortical neurons were used in similar experiments to compare the results obtained with DA neurons to a population of neurons known to establish mostly synaptic-type terminals. We found that NL-1^AB^ also induced a robust recruitment of cortical terminals (**Fig 7F**) as previously reported (6). Quantification of the total intensity of VGluT1 puncta on HEK293T cells expressing NL-1^AB^ showed an 18-fold increase compared to the control CD4 condition (**Fig. 7G**). These results suggest that NL-1^AB^ has efficient synaptogenic activity to induce presynaptic differentiation of DA neurons, to a level comparable to that observed for cortical neurons. This observation suggests that the limiting factor preventing most of the numerous terminals established by DA neurons to form synapses is more likely to be at the presynaptic rather than at the postsynaptic level.

**Figure 7.**
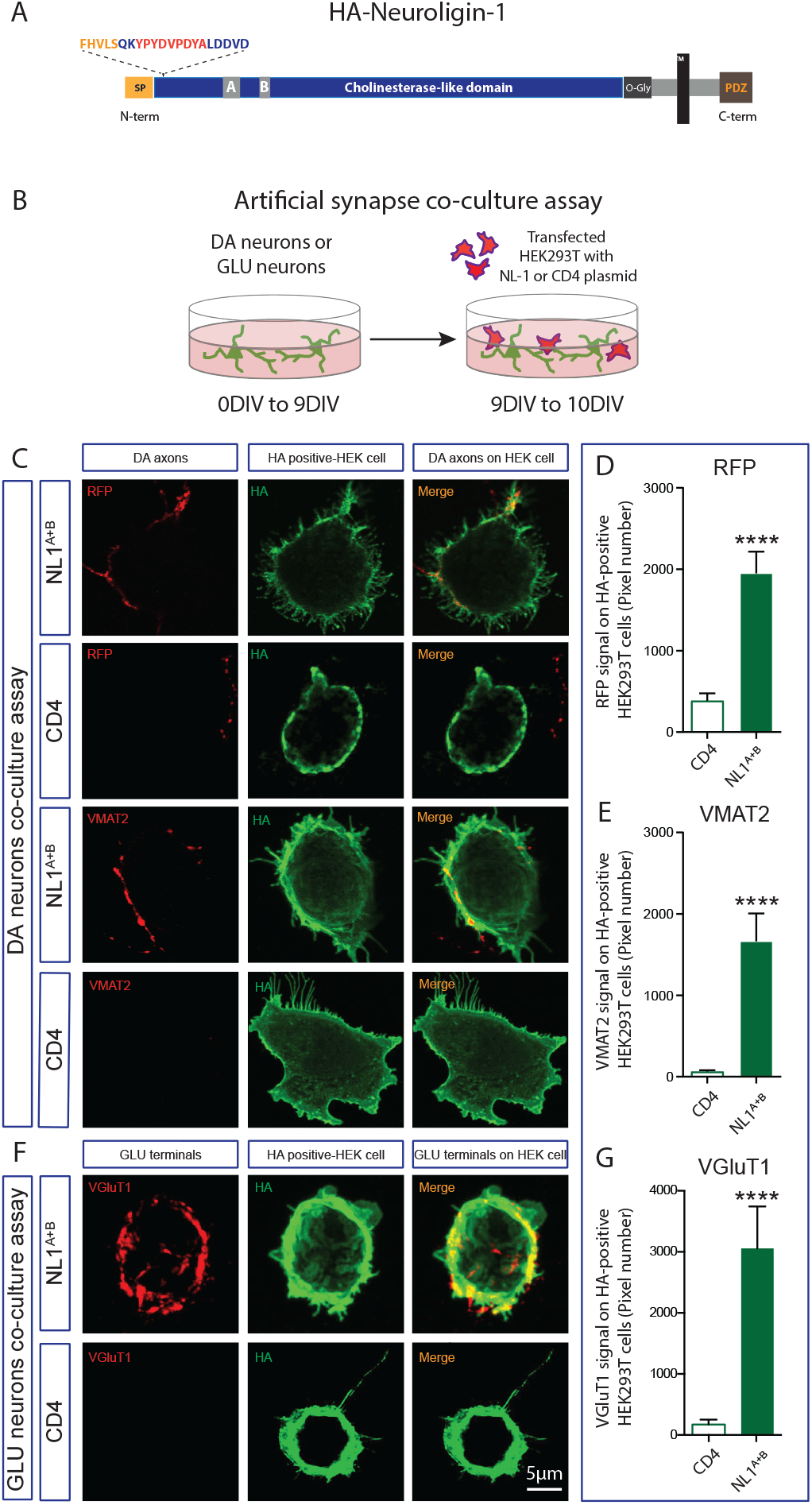
NL-1 induce presynaptic differentiation of DA neuron terminals. **A-** Schematic representation of NL-1-HA, epitope-tagged in the extracellular domain**. B-** Schematic representation of the neuron-HEK393T cell co-culture assay**. C-** Representative images of double immunolabelling for RFP (red) or VMAT2 (red), identifying terminals from DA neurons and HA-tagged (green) NL-1 construct expressed by co-cultured HEK293T cells. HA-CD4 was used as a negative control protein due to its known lack of synaptogenic activity. **D & E -** Quantification of total intensity of RFP (**D**) and VMAT2 (**E**) axonal puncta on HEK293T cells expressing NL-1-HA or CD4-HA. **F**- Representative images of double immunolabelling for VGluT1 (red), identifying cortical glutamate terminals and HA-tagged (green) NL-1 construct expressed in co-cultured HEK293T cells. HA-CD4 was used as a negative control. **G -** Quantification of total intensity of VGluT1 axonal puncta on HEK293T cells expressing NL-1-HA or CD4-HA. Bar graphs represent the mean + SEM, t-test, ^*\*\*\**^*p<0.001;* ^*\*\*\*\**^*p<0.0001*. HEK cells=23 (NL-1) and HEK cells= 25 (CD4) from 2 different experiment with glutamatergic neurons; HEK cells=58 (NL-1) and HEK cells=49 (CD4); RFP group from 2 different experiments with DA neurons; HEK cells=25 (NL-1) and HEK cells=25 (CD4); VMAT2 group from 2 different experiments with DA neurons. The number of observations represents the number of fields from individual HEK293T cell examined.

### *Overexpression of Nrxn 1*α^*SS4-*^ promotes the formation of both excitatory and inhibitory synapses by DA neurons

One of the most important transsynaptic binding partners of NL-1 is the presynaptic protein Nrxn-1. This protein is also involved in glutamatergic synapse formation and has two versions: the long version, α-neurexin (Nrxn 1α), and the short version, ß-neurexin (Nrxn 1ß). Both Nrxn 1α and Nrxn 1ß, regulate excitatory synapse formation and function (30, 57). Considering that a subset of DA neurons uses glutamate as a second neurotransmitter, we first investigated the endogenous expression of Nrxn 1 in DA neurons. We found that both isoforms are expressed at high level in both VTA and SNc DA neurons, although Nrxn 1α was expressed at higher levels in VTA DA neurons (**Fig. S6**). We therefore examined whether overexpression of Nrxn 1α influences synapse formation by these neurons. Lentiviral overexpression of Nrxn 1α (splice version SS4-; Nrxn 1α^SS4-^) in DA neuron co-cultures (**Fig. 8A** and **8B**) caused a 2-fold increase in the proportion of DA terminals colocalizing with the excitatory postsynaptic organizer PSD95 compared to the control group (**Fig. 8C** and **8D**). The average size of PSD95 positive synaptic puncta or their total area was unchanged (**Fig. 8E** and **8F**), suggesting that the overexpression of Nrxn1α^SS4-^in DA neurons triggered the establishment of more contacts with pre-existing postsynaptic clusters. Overexpression of Nrxn1α^SS4-^in DA neurons induced a similar increase in the proportion of DA terminals colocalizing with the inhibitory postsynaptic protein Gephyrin (**Fig. 8G** and **8H**), with no change in Gephyrin puncta size or total Gephyrin puncta area (**Fig. 8I** and **8J**).

**Figure 8.**
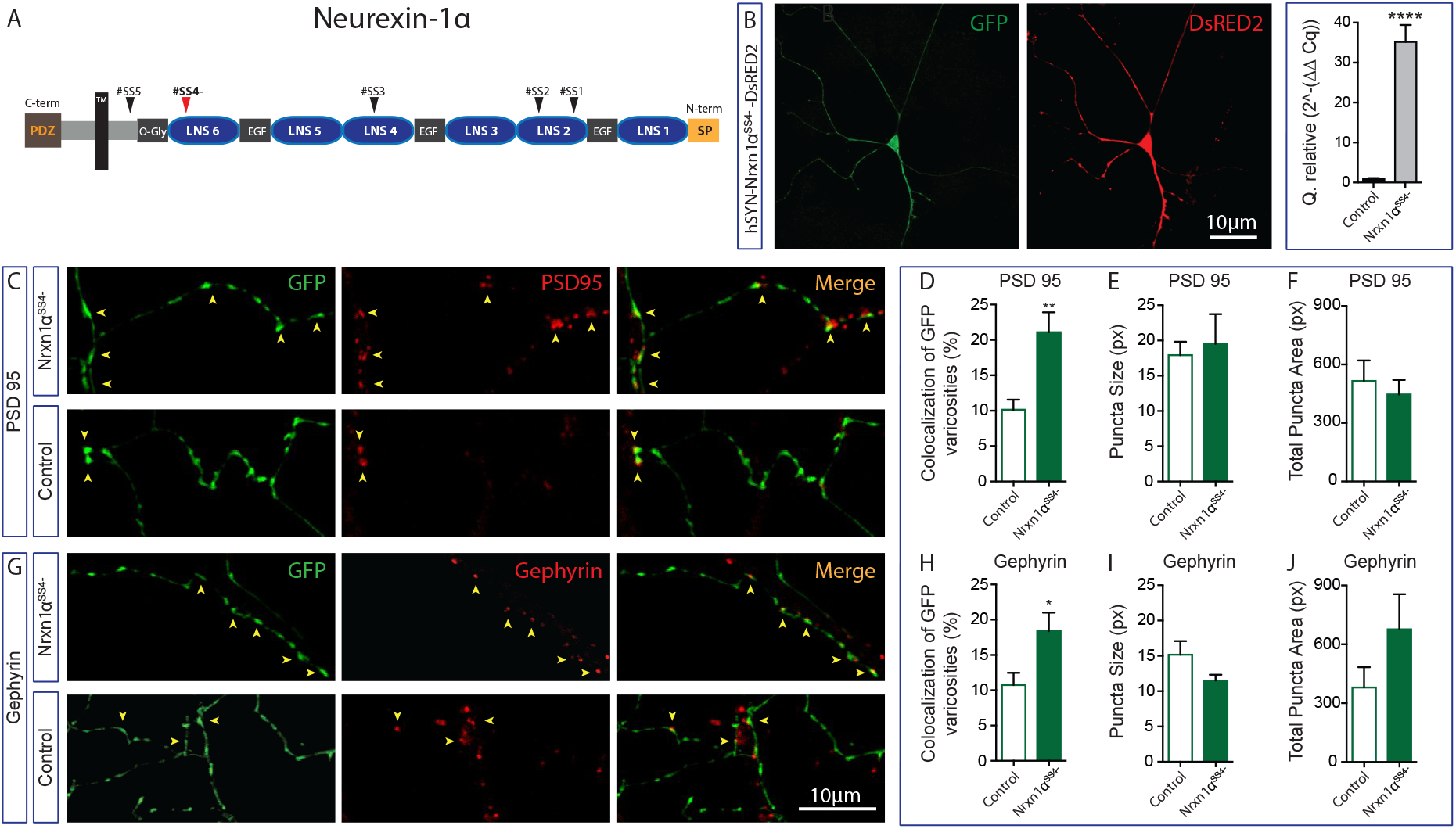
Overexpression of Nrxn 1α^SS4-^promotes the formation of both excitatory and inhibitory synapses by DA neurons. **A**- Schematic representation of the Nrxn 1α^SS4**-**^without the splicing site 4 (SS4-). **B**- Immunofluorescence images showing expression of DsRed2 expressed together with Nrxn 1α^SS4**-**^in a GFP-positive DA neuron. The bar graph shows a relative quantification by qPCR of mRNA encoding for Nrxn 1α^SS4**-**^after neuronal transfection with the plasmid pLV-hSYN-**Nrxn-1α**-DsRED2 compared to the control plasmid pPL-hSYN-DsRED2. **C**- Representative images of double immunolabelling for GFP, expressed by DA neurons and PSD95 after overexpression of Nrxn 1α^SS4-^or of the control construct. **D** to **F -** Summary graphs representing the proportion of excitatory synapses established by DA neurons (**D**), including the average puncta size (**E**) and the total puncta area (**F**) of PSD95 signal. **G**- Representative images of double immunolabelling for GFP, expressed by DA neurons and Gephyrin after overexpression of Nrxn 1α^SS4-^or of the control construct. **H** to **J -** Summary graphs representing the proportion of excitatory synapses established by DA neurons (**H**), including the puncta size (**I**) and the total puncta area (**J**) of Gephyrin signal. Bar graphs represent the mean + SEM, t-test, Tukey’s multiple comparison test, ^*\*\*\**^*p<0.001;* ^*\*\*\*\**^*p<0.0001*. DA neurons n=31 and n=41 (pLV-hSYN-Nrxn1α^SS4-^-DsRED2 and pLV-DsRED2 respectively; PSD95 group) and DA neurons n=27 and n=26 (pLV-hSYN-Nrxn1α^SS4-^-DsRED2 and pLV-DsRED2 respectively; Gephyrin group) from 3 different cultures. The number of observations represents the number of fields from individual transfected neurons examined.

## DISCUSSION

In this study we provide a novel perspective on the connectivity of DA neurons with an exhaustive characterization of the axonal domain of this key neuromodulatory brain system. We found that DA neurons establish a highly distinctive axonal domain with a majority of non-synaptic terminals, in a way that is fundamentally different compared to cortical glutamatergic and striatal GABAergic neurons. Both synaptic and non-synaptic DAergic varicosities express basic presynaptic proteins such as Syt1, suggesting that the majority of these varicosities have the machinery to release neurotransmitters. We discovered that synaptic and non-synaptic terminals differ in their expression of active zone structural proteins: the active zone protein bassoon was found to be mainly expressed in DA terminals located close to a target cell, while most non-synaptic terminals were devoid of bassoon. The active zone protein RIM1/2 was more sparsely expressed in both synaptic and non-synaptic DA terminals. Using the activity-dependent synaptic vesicle probe FM1-43, we found that the majority of DA terminals, both synaptic and non-synaptic, are active. Finally, providing initial insight into the mechanistic underpinnings of this unique connectivity, we find that the postsynaptic trans-synaptic protein NL-1 is able to efficiently trigger formation of synapse-like structures by primary DA neurons, while overexpression of the presynaptic trans-synaptic protein Nrxn-1 causes an increase in the proportion of terminals adopting a synaptic configuration.

### Heterogeneity in DA release sites

We found that ~80% of axonal-like varicosities established by DA neurons express the exocytosis calcium sensor Syt1, suggesting that the SNARE complex and associated proteins are present in the majority of axonal varicosities established under our experimental conditions (**Fig. 2A** and **2F**). Although we have not examined in detail the presence of other vesicular exocytosis proteins in the present study, most of these varicosities also contain SNAP25 and SV2. The presence of Syt1 in terminals has previously been shown to provide a good index of functionality (1). Perhaps a bit surprising, the expression of VMAT2, considered as a specific marker of terminals that release DA or other monoamines was found in only ~55% of DA terminals (**Fig. 2B** and **2G**). These results suggest that only half of DA terminals can package and release DA. One possible explanation of this low proportion is the fact that some DA neurons are also able to package and co-release other neurotransmitters at some of the axon terminals, including glutamate and GABA (11, 14, 50, 58). Furthermore, recent studies have reported that co-release of DA and glutamate mainly occurs from separate, segregated sets of terminals (52, 59). The present experiments were performed with primary DA neurons in co-culture with striatal neurons. The medial part of the VTA and the lateral part of the SNc were used. Because previous work reported that VMAT2 is more expressed in DA neurons located in the lateral part of the VTA compared to the medial part (60), it is possible that this may have contributed to our finding of a large proportion of terminals without detectable VMAT2. Also, recent work suggested that 75% of DA neurons in the mesencephalon express the mRNA encoding for VMAT2 suggesting that a small subset doesn’t express VMAT2, at least at this time point *in vitro* (52). Additional studies will be required to examine this further and to determine whether this phenomenon is developmentally regulated.

### The active zone proteins bassoon and RIM are mainly segregated

Our work provides new insights on the differential molecular make up of synaptic and non-synaptic DA terminals. At classical synapses, release sites are characterized by the presence of active zones that contain different scaffold proteins including RIM, RIM-BP, bassoon/piccolo or ELKS (48, 61). The active zone protein RIM has been identified as a key scaffold protein notably involved in vesicle priming and Ca^2+^ channel tethering (3, 62). Bassoon is also an important scaffold protein and plays a role in synaptic vesicle clustering without participating directly in synaptic vesicle exocytosis (63). In our *in vitro* system, we first found that only ~30% of DA terminals contained bassoon. Strikingly, we observed that the majority of them were in close proximity or in contact with a target cell (**see Fig. 4** and **S3**). We also found that only ~30% of bassoon positive DA terminals were also positive for RIM1/2 (**Fig. S4F**). Conversely, only a third of RIM1/2 positive terminals contained bassoon (**Fig. S4E**). The mechanism and functional implications of this differential expression of active zone proteins in DA terminals are presently unclear. Interestingly, the conditional KO of RIM1 and RIM2 in DA neurons was recently shown to completely block evoked DA release, while extracellular levels of DA were only partially decreased (49). This finding suggests the possibility that while action potential-evoked DA release requires RIM1/2, spontaneous release occurs through a separate mechanism. Spontaneous DA release could thus potentially be mediated by Syt1- and SNARE-dependent exocytosis without any implication of active zone proteins, as previously suggested for glutamate release from cortical neurons (64). In our work, we found that approximately 80% of DA terminals contain Syt1, suggesting that most of them have the capacity to release DA, spontaneously or in an activity-dependent manner. Our ultrastructural observations with TEM also support this conclusion as most varicosities examined contained large pools of synaptic vesicles (**Fig. 1L**). Our observation of the selective presence of the active zone scaffold protein bassoon at DA terminals located in close proximity to a target cell is particularly intriguing. Previous work in hippocampus has suggested that target cells can play a role in synapse maturation during synaptogenesis and that some neurons also possess non-synaptic release sites that are functional and mobile and that express the active zone protein bassoon (65, 66). Although we hypothesize that target-derived signals are likely to be required to restrict bassoon to release sites that are in close proximity to target cells, further work will be required to test this directly.

### DA neurons establish a minority of excitatory, inhibitory and pure DA synapses

Our experiments provide a new perspective on the synaptic and non-synaptic terminals established by DA neurons. Synapses are classically defined by an apposition between a pre and a post synaptic domain (67, 68). We investigated if the GFP-positive presynaptic terminals were associated with a postsynaptic specialization. Based on previous *in vivo* ultrastructural observations suggesting that only a small subset of DA terminals is associated with a postsynaptic site, we first used MAP2 as a marker of potential post-synaptic elements. We found that only ~20% of VTA and SNc DA terminals are in close proximity to MAP2-positive dendritic domains (**Fig. 2E** and **2H**). These data are consistent with previous studies suggesting that DA neurons, *in vivo* and *in vitro*, are able to develop terminals without showing any clear apposition to a postsynaptic site (43, 69, 70). The molecular mechanism allowing DA neurons to differentially establish synaptic and non-synaptic terminals is presently unknown. Here again, one hypothesis to consider is that target cells provide signals that retrogradely determine the structure of terminals that become synaptic, while the non-synaptic terminals would possibly develop along a default non-synaptic differentiation path. In accordance with this hypothesis, recent work has revealed how striatal target cells regulate the glutamatergic phenotype of a subset of terminals established by DA neurons (52).

While glutamatergic synapses established by DA neurons in the ventral striatum of rats have been described to possess a synaptic organization (71, 72), the possibility that DA neurons establish terminals that form synapses containing DA receptors has been previously considered, although supported by very limited evidence. Previous work using electron microscopy has indeed suggested that dopaminergic varicosities can be found associated with postsynaptic specializations that contain D1 receptors (24). In the present work, we took advantage of a newly generated genetically modified mouse model allowing to easily visualize DA D2 receptors without the important limitation due to the limited specificity of antibodies directed against DA receptors. Our findings reveal a small proportion of DAergic terminals that colocalize with postsynaptic D2 receptor clusters. These results demonstrate that under our experimental conditions, only a very small subset of less than 10% of DAergic terminals form what could potentially be “pure” DA synapses. Taken together, our results suggest an intrinsic ability of DA neurons to preferentially develop a majority of non-synaptic terminals. We hypothesize that while the small subset of synaptic release sites provides different types of fast signals to target cells, mediated through DA, glutamate and GABA receptors, the larger subset of non-synaptic terminals represents the morphological substrate of slower signals previously described as “volume” or “diffuse” transmission (20, 22, 70, 73–75).

### A majority of DA terminals are active

While DAergic neurons are endowed with an exceptionally arborized axonal compartment, with a very large number of axonal-like varicosities, whether all or a subset of these varicosities represent functional release sites in unclear. Interestingly, previous experiments performed with a newly developed DA false transmitter up taken by VMAT2, suggested that only ∼20% of DAergic varicosities are able to release DA (76). Here we took advantage of the high signal to noise ratio and simplicity of our *in vitro* model to examine whether the large numbers of varicosities established by DA neurons are active. We chose to evaluate this using a FM1-43 uptake assay, an approach that labels activity-dependent, actively recycling vesicles, irrespectively of their neurochemical identity. We found that a large majority of axonal varicosities established by DA neurons are active, suggesting that the majority of non-synaptic terminals represent functional neurotransmitter release sites, even if they are not associated with a tightly linked postsynaptic specialization. The different conclusions of the present study compared to that of Pereira and colleagues could result from several technical considerations. One of these is that this earlier study used a false fluorescent neurotransmitter thought to selectively label DAergic terminals containing VMAT2, while in the present study, we examined terminals irrespective of whether they contained VMAT2, VGluT2 or potentially other neurotransmitters such as peptides. This could have potentially provided a broader overview of all terminals, especially considering that we found a large subset of syt1-positive terminals established by DA neurons but devoid of detectable VMAT2 (**see Fig.2G** and **Fig.5G**). Compatible with our observation, a previous study reported that the majority of axonal varicosities established by DA neurons have the capacity of load and release FM1-43 (69). Finally, we found that synaptic and non-synaptic terminals exhibited similar FM1-43 destaining properties, suggesting that both types possess similar activity-dependent mechanisms that subsequently mediate fast and slow signals.

### Presynaptic Nrxn-1α^SS4-^ and postsynaptic NL-1^AB^ influence synapse formation by DA neurons

Using an artificial synapse co-culture assay and a lentiviral vector to overexpress a transsynaptic protein on primary DA neurons, in this study we found that the postsynaptic protein NL-1^AB^ expressed on HEK293T represents an efficient signal to induce presynaptic differentiation of DAergic terminals. We also found that presynaptic Nrxn1α^SS4**-**^overexpression increases the number of excitatory and inhibitory synapses established by DA neurons.

The NL-1 protein is preferentially found at excitatory synapses and directly interacts with the scaffolding protein PSD95 (55, 77, 78). The maintenance and the function of excitatory synapses are regulated by NL-1 via the recruitment of AMPA and NMDA receptors at the postsynaptic domain (55). In our *in vitro* experiments, we clearly demonstrated that HEK293T cells expressing the major form of NL-1 (splice variant A^+^B^+^; NL1^AB^) attract DA terminals, suggesting that NL-1^AB^ has the potential to induce presynaptic differentiation of axon DA terminals. An implication of NL-1 in the establishment of synapses by DA neurons is in line with the fact that some of these neurons establish glutamate synapses and release glutamate as a second neurotransmitter at sites that are synaptic in structure (22). This result is also compatible with our observation that HEK293T cells expressing NL-1^AB^ are able to induce the formation of artificial synapses by DA neuron terminals and with our finding of robust expression of Nrxn-1ß by DA neurons (**Fig. 7** and **Fig. S6**). Nrxn-1ß strongly interacts with NL-1 containing the splice site B, whereas Nrxn-1α is not able to (30).

Considering this ability of NL-1 to promote synapse formation by DA neurons, it is puzzling that in the intact brain, expression of NL-1 by striatal neurons appears to promote glutamate synapse formation by cortical neurons but not by DA neurons. Further experiments will be needed to solve this important question and to determine which others pre- and postsynaptic proteins regulate synapse formation by DA neurons. It is interesting to note that recent work has provided evidence supporting the possibility that NL-2 regulates the formation of synapses by DA neurons (8, 79). One possibility is that NL-2 plays a role in the establishment of GABA release sites by DA neurons (10, 50).

In the context of the strong ability if NL-1 to promote synapse formation by DA neurons, a limited formation of synapses by DA neurons *in vitro* and *in vivo* could perhaps be due to limited levels of Nrxn-1 at individual terminals. Although we found that Nrxn-1α and Nrxn-1ß are robustly expressed by DA neurons, the fact that DA neurons in rodents are thought to establish upwards of 100 000 individual release sites along their axonal arbor (15) could give rise to limited levels of presynaptic transsynaptic proteins in these terminals. Nrxn-1 levels could therefore represent a limiting factor in the establishment of synapses by DA neurons. Our finding that lentiviral overexpression of Nrxn-1α (splice variant SS4-; Nrxn-1α^SS4-^) in DA neurons leads to an increase in the proportion of terminals establishing synapses agrees with this hypothesis. Further experiments will be needed to determine if these newly created synapses are functional and to evaluate whether other Nrxn isoforms and other transsynaptic proteins can also influence the synaptic connectivity of DA neurons *in vitro* and in the intact brain.

Together, our findings shed new light on the peculiar capacity of DA neurons to establish a highly complex axonal arbor endowed with a large repertoire of signaling mechanisms. We envision that the striking, seemingly default capacity of DA neurons to establish an extensive axonal arbor endowed with a large numbers of axon terminals that appear to actively avoid direct contact with target cells is likely to be due to a developmental program that is shared with other key modulatory neurons including those using serotonin, norepinephrine and acetylcholine. Further work will be required to determine the common mechanisms involved.

## Acknowledgments and funding’s

The authors wish to thank Brooks Robinson and John T. Williams (Oregon Health and Science University) for providing the D2(GFP) knock-in mouse line and for their comments on an early version of this manuscript. We also thank Guillaume Fortin for his help in some of the experiments. This work was funded by a grant from Canadian Institutes of Health Research (CIHR; MOP106556) to L-E Trudeau. Charles Ducrot received salary support from the GRSNC, Parkinson Canada and from the Fonds du Québec en Recherche, Santé (FRQS). Martin Parent and Constantin Delmas were funded by a grant from CIHR (MOP153068). Ulrich Gether, Freja Herborg and Matthew Domenic Lycas were funded by the Lundbeck Foundation (2018-792; 2017-4331 and 2016-3154). Hideto Takahashi was funded by a grant from CIHR (PJT159588) and received salary support though the FRQS Research Scholar (Senior) program. Aurelie Fallon received financial support from CIHR, the department of neurosciences of the Université de Montréal, and FRQS in collaboration with Parkinson Quebec. SEM analyses were carried out in the Electron imaging Facility of Université de Montréal. Dainelys Guadarrama received salary support from the FRQS.

## Author contributions

Charles Ducrot designed, performed, and analyzed the majority of experiments related to synapse quantifications. Marie-Josée Bourque and Charles Ducrot prepared all cell cultures. Charles Ducrot performed and analyzed all the experiments related to FM1-43. Anne-sophie Racine performed all experiments related to the olfactory bulb and participated to some of the synapse quantifications. Charles Ducrot and Marie-Josée Bourque prepared the cultures and resin inclusions for TEM experiments. Constantin Delmas performed all the sections, observations and analyses related to the TEM experiments. Matthew Domenic Lycas and Benoît Delignat-Lavaud performed the dSTORM experiments. Matthew Domenic Lycas analyzed all the data related to the dSTORM experiments. Charlotte Michaud-Tardif performed some of the experiments related to the presynaptic characterization. Samuel Burke Nanni performed the analysis related to the density of DA terminals. Charles Ducrot and Dainelys Guadamarra Bello performed the experiments related to the SEM. Charles Ducrot and Aurélie Fallon performed the artificial synapse experiments. Antonio Nanci provided guidance for SEM sample processing. Louis-Eric Trudeau designed and coordinated the project and contributed to writing of the manuscript. Martin Parent, Hideto Takahashi, Antonio Nanci, Freja Herborg and Ulrik Gether provided conceptual feedback and revised the manuscript.

