## Supplementary material for "Dopaminergic neurons establish a distinctive axonal arbor with a majority of non-synaptic terminals": Fig. S

### Fig.S1

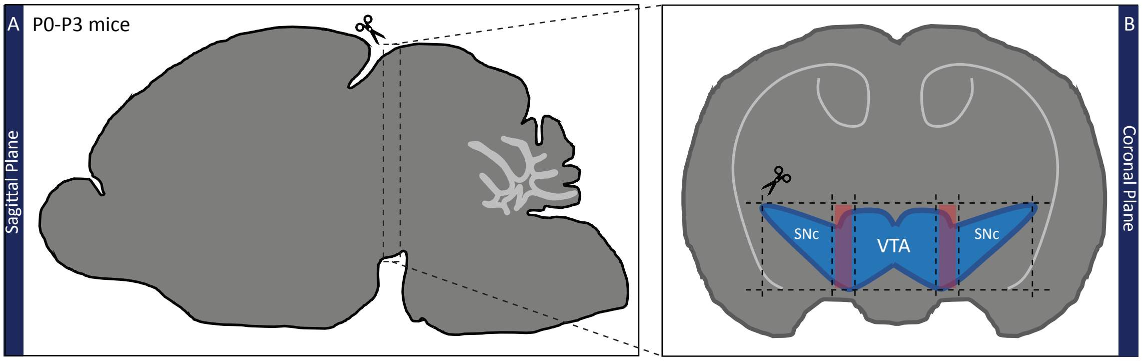

### Fig. S2

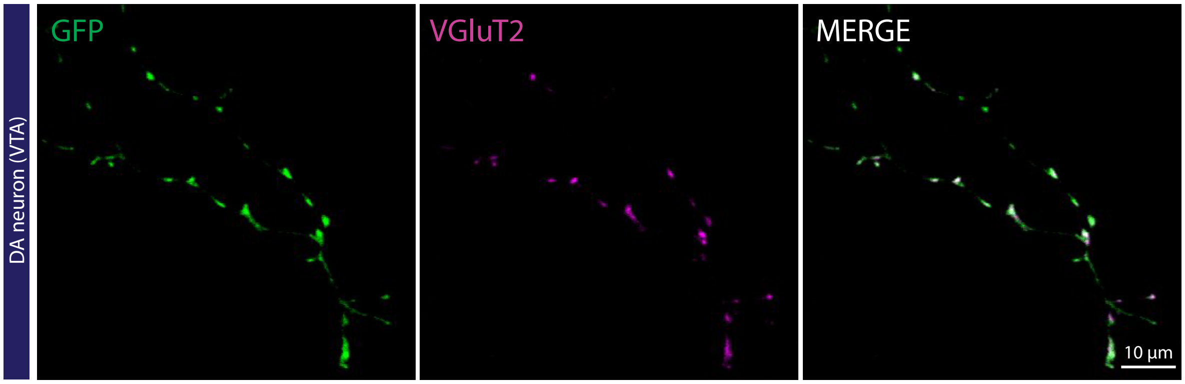

### Fig. S3

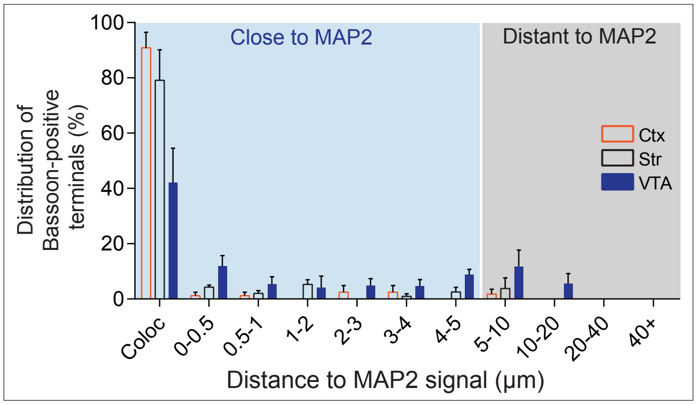

Close to MAP2
Distant to MAP2
Distribution of Bassoon-positive terminals (%)
100
80
60
40
20
0
Ctx
Str
VTA
Coloc
0-0.5
0.5-1
1-2
2-3
3-4
4-5
5-10
10-20
20-40
40+
Distance to MAP2 signal (µm)

### Fig. S4

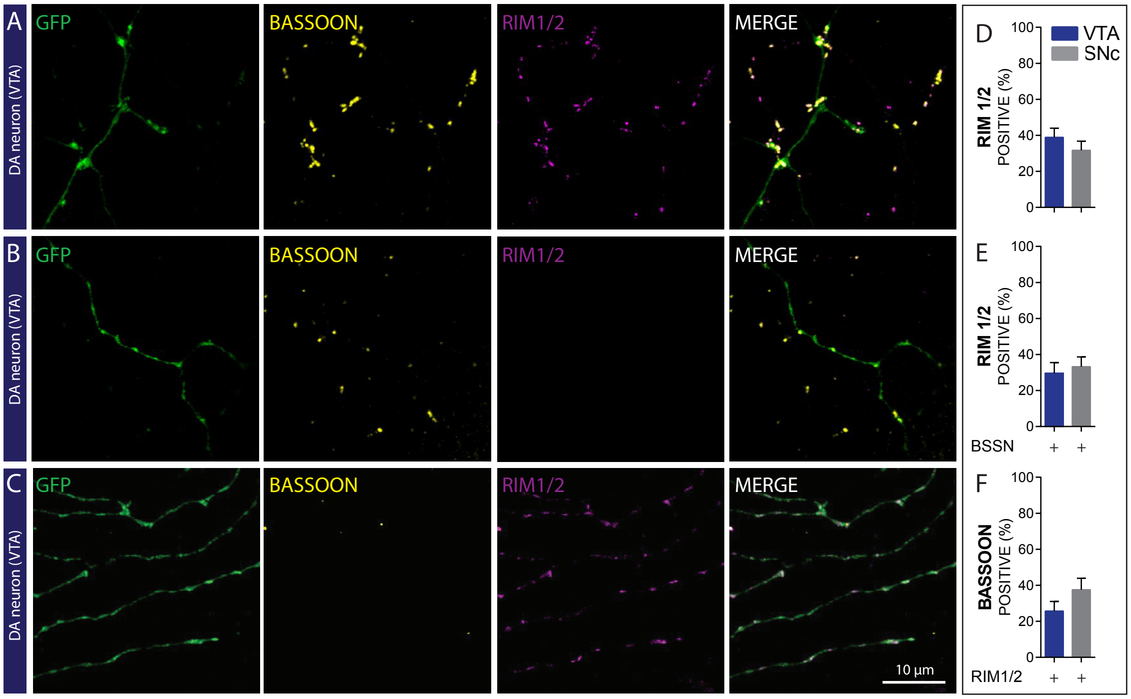
A
B
C
DA neuron (VTA)
GFP
BASSOON
RIM1/2
MERGE
10 μm
D
E
F
VTA
SNc
RIM 1/2
POSITIVE (%)
BASSOON
BSSN
RIM1/2
+
0
20
40
60
80
100

### Fig. S5

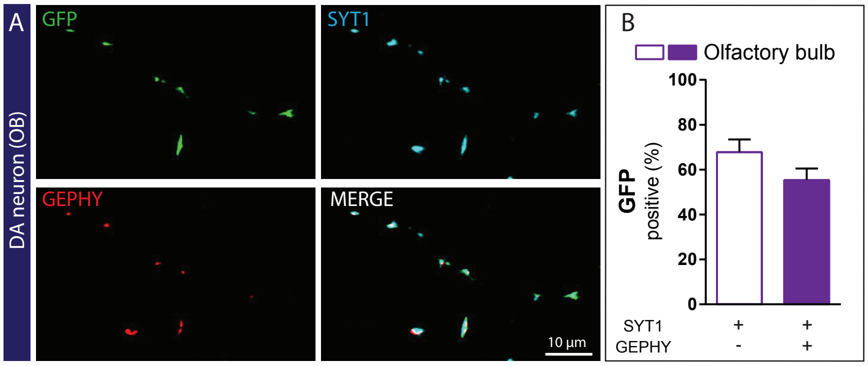

A
DA neuron (OB)
GFP
SYT1
GEPHY
MERGE
10 µm
B
Olfactory bulb
GFP
positive (%)
100
80
60
40
20
0
SYT1
GEPHY
+
+
-
+

### Fig.S6

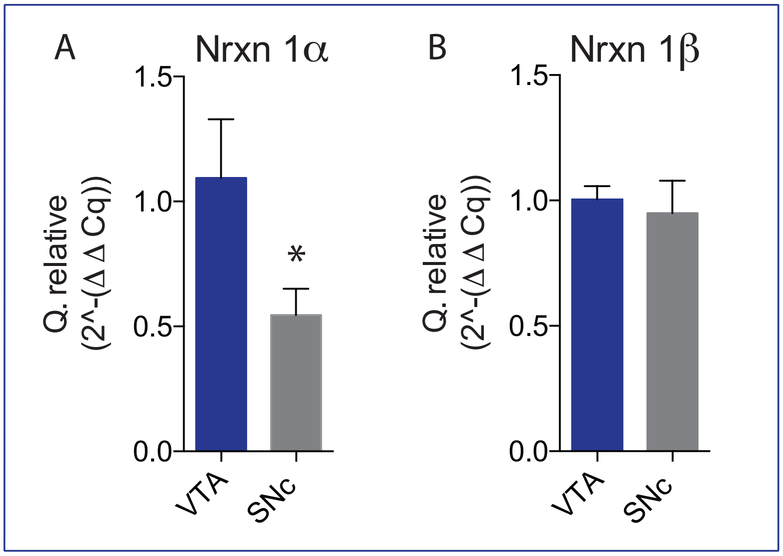

A
Nrxn 1α
Q. relative
(2^-(Δ Δ Cq))
1.5
1.0
0.5
0.0
*
VTA
SNc
B
Nrxn 1β
Q. relative
(2^-(Δ Δ Cq))
1.5
1.0
0.5
0.0
VTA
SNc

#### SUPPLEMENTAL FIGURE LEGENDS

**Supplemental Fig. S1 - Schematic representation of the forebrain and midbrain dissection**. **A-** The forebrain was first isolated by dissection along the dashed line. **B-** Schematic representation of a coronal view of the dissected forebrain. The dashed lines represent the parts used to prepare the primary cultures of VTA and SNc DA neurons. The red squares represent parts that were excluded.

**Supplemental Fig. S2 – DA neurons express VGlut2 at some of the axon terminals.** Photomicrographs illustrating GFP positives axonal varicosities along the axonal arborization of a DA neuron expressing VGluT2.

**Supplemental Fig. S3 – Bassoon distribution along the axonal arbor of DA neurons compared to glutamatergic and GABAergic neurons.** Histograms representing the mean +/- SEM from 3 different fields from 3 different neurons for each structure (VTA, cortex (Ctx) and striatum (Str)). Each histogram includes 137 terminals from cortical neurons, 134 terminals from striatal neurons and 612 terminals from VTA DA neurons.

**Supplemental Fig. S4 – The active zone proteins RIM 1/2 and bassoon are partially segregated in distinct sets of DA terminals. A-** Photomicrographs illustrating GFP positives axonal varicosities established by VTA DA neurons expressing bassoon and co-expressing RIM1/2. **B-** Photomicrographs illustrating GFP positives axonal varicosities established by VTA DA neurons expressing bassoon but not RIM1/2. **C**- Photomicrographs illustrating GFP positives axonal varicosities established by VTA DA neuron expressing RIM1/2 but not bassoon. **D**- Bar graph representing the proportion (%) of VTA and SNc GFP positive axonal varicosities expressing RIM1/2. **E**- Bar graph representing the proportion (%) of DAergic GFP positive axonal varicosities positive for bassoon and colocalizing with RIM1/2. **F**- Bar graph representing the proportion (%) of DAergic GFP positive axonal varicosities positive for RIM1/2 and colocalizing with bassoon. The bar graphs represent the mean + SEM. VTA n=28 SNc n=30; from 2 different cultures. The number of observations represents the number of fields from individual neurons examined

**Supplemental Fig. S5 – Olfactory Bulb DA neurons establish terminals that are mainly synaptic.** **A –** Photomicrographs illustrating GFP positives varicosities along the axonal arborization of an olfactory bulb DA neuron expressing Syt1 and colocalizing with gephyrin. **B –** Bar graph representing the proportion (mean +/- SEM) of GFP/Syt1 positive axonal varicosities colocalizing with gephyrin (purple) or not colocalizing with gephyrin (white), established by olfactory bulb DA neurons. VTA n=27 SNc n=27; from 3 different cultures. The number of observations represents the number of fields from individual neurons examined.

**Supplemental Fig. S6 - Nrxn-1α and Nrxn-1ß genes are expressed in DA neurons.** Relative quantification by qPCR of mRNA encoding for Nrxn-1α (**A**) and Nrxn-1ß (**B**), for both VTA and SNc DA neurons.
